# Asymmetric Costs of Coordination Errors Generate Selection on Social and Sensorimotor Phenotypes in an Animal Collective

**DOI:** 10.64898/2026.07.09.737300

**Authors:** Luke C. Larter, Michael J. Ryan, Matthew J. Fuxjager

**Affiliations:** Institute at Brown for the Environment and Society, Brown University, 85 Waterman St, Providence, RI 02912, USA; Department of Ecology, Evolution, and Organismal Biology, Brown University, Providence, RI 02912, USA; Center for Animal Behavior, Smithsonian Tropical Research Institute, Apartado 0843-03092, Balboa, Republic of Panama; Department of Integrative Biology, University of Texas at Austin, 2415 Speedway, Austin, TX 78712, USA

## Abstract

Animal collectives are dynamic social environments where individuals must continually coordinate their behavior with neighbors. Sensorimotor limitations like processing delays make coordination errors inevitable and, in high-stakes contexts like collective predator evasion and courtship, errors can be extremely costly. Identifying errors and quantifying their costs are challenging in flocks and schools, so collective behavior models traditionally treat errors as random noise. However, if instead the rate and severity of costly errors within collectives are biased towards certain phenotypes, selection against error-prone phenotypes is a neglected feature of the selective landscape. We tested this in experimental choruses of a call-alternating frog (túngara frogs, *Physalaemus* [*=Engystomops*] *pustulosus*). Here inadvertent synchronous calls represent discrete coordination errors attributable to sensorimotor limitations rather than strategy, and they carry steep attractiveness costs; mate-searching females disfavor synchronous calls, especially the following calls of synchronous call pairs. Synchronous calls were frequent, and synchrony outcomes varied non-randomly by social and sensorimotor phenotypes. Inter-male compatibility in intrinsic call rhythms structured synchrony, causing males with slower rhythms to synchronize at higher rates. Furthermore, during synchrony, males with longer response latencies more often produced costlier following calls. Bayesian simulations combining interaction data with experimentally derived female preference estimates revealed that these mechanistic linkages between male phenotypes and synchrony outcomes generated systematic female biases against males with slower call rhythms and longer response latencies. Thus, coordination errors are not unstructured noise. Rather, phenotype-dependent variation in the frequency and/or severity of errors can generate directional selection on social phenotypes and their sensorimotor underpinnings in animal collectives.

## Introduction

In animal collectives, individuals coordinate their behavior with neighbors to navigate high-stakes ecological contexts like predator evasion and collective courtship [*1,2,3*]. Here, mistimed or poorly calibrated responses can result in death or missed mating opportunities, which selects for precise coordination strategies [*3,4,5,6*]. However, collectives are fast-paced and dynamic, and it takes time to process sensory information received from neighbors, then select and mount a motor response [*7,8*]. Consequently, the split-second decisions that underpin ongoing coordination must be based on stale information, or imperfect extrapolations derived from it [*9,10*]. This suggests that coordination errors, *i.e.* discrepancies between target and realized coordination outcomes, will remain common despite their potentially high costs.

Coordination errors are an emergent social phenomenon; they arise not just from sensorimotor limitations, but from how those limitations manifest themselves when a given social strategy is pursued in a given social context. Individuals in single-species collectives possess qualitatively similar limitations [*11*] and follow qualitatively similar social strategies [*6*]. This likely biases errors away from being purely random, resulting in the same structured error types arising repeatedly. When errors exhibit consistent structure, even subtle inter-individual variation [*12*] in the capacity to minimize the rate and severity of common error types could yield robust asymmetries in accumulated error-derived costs or risk exposure. This would then generate selection gradients that favor phenotypes that are more effective at mitigating the costs of common error types. As noted, errors emerge as sensorimotor constraints interact with expressed social strategies, meaning selection for error mitigation could proceed by refining sensorimotor or social phenotypes, or both synergistically. However, despite their presumed selective importance, empirical studies linking variation in sensorimotor or social phenotypes to error propensities and downstream selection are lacking.

One reason the selective effects of coordination errors remain obscure is that studying them in locomotory collectives like bird flocks and fish schools is immensely difficult. Here, coordination takes the form of incremental alignment of heading and speed, and inter-neighbor spacing. Categorical errors such as collisions are rare, and definitively identifying errors of degree is challenging. Thus, though recent empirical work incorporates varied perceptual [*13,14*] and locomotor limitations [*15,16*] when modeling interaction outcomes, these principles are seldom applied explicitly to coordination errors. Furthermore, traditional collective behavior models represent errors as random noise terms applied uniformly across individuals [*17,18*]. However, if variation in error mitigation abilities causes error-derived costs to be distributed asymmetrically, models omitting this do not fully capture the selective landscape honing social phenotypes and their sensorimotor foundations.

Addressing these gaps requires a system where coordination errors are discrete and unambiguous, such that individualized error rates and outcomes can be quantified, and their drivers investigated empirically. Acoustic collective behavior in chorusing frogs meets these criteria. Various frogs form choruses in which males call to attract females to mate [*19*]. In many species, call overlap can disrupt salient call properties and makes calls difficult for females to recognize or locate [*20*]. Thus, overlapping calls are disfavored by females relative to unobscured calls [*21*]. This selects for sensorimotor coordination algorithms that allow males to alternate their calls with those of nearby rivals without overlap [*3,22*]. Yet, overlap remains prevalent, especially in crowded choruses. Call overlap in primarily alternating frogs comes in various forms within and across taxa, some of which are actively facilitated in certain contexts as part of flexible social strategies [*23,24*]. Thus, not all forms of call overlap can be assumed to be coordination errors born of sensorimotor constraints.

However, one specific form of overlap is ubiquitous across alternating frogs, and its origins as a sensorimotor coordination error are unequivocal: synchrony. During synchrony the onsets of two or more males’ calls occur near-simultaneously [*3,25,26*], and the timing of synchronous call onsets bears a diagnostic signature of the sensorimotor limitation that led to synchrony. The ‘effector delay’ [*sensu 3*] is the intrinsic latency between a call being triggered by the nervous system and produced by the vocal apparatus (*i.e.* response latency), and it generates synchrony in the following way. Ordinarily, males are inhibited from initiating calls while perceiving a rival’s call, which facilitates call alternation without overlap. However, when a rival’s call begins during a male’s effector delay, this rival’s call cannot inhibit initiation of this male’s already impending call [*3*]. Consequently, both individuals’ calls are initiated near-simultaneously. This mechanistic link between effector delays and synchrony can be verified empirically, by the fact that delays between synchronous call onsets are always less than or equal to the limit of that species’ effector delay (*26*–*28*; Fig. 1E). Synchrony is therefore highly stereotyped (Fig. 1C) and clearly identifiable as an incidental failure of alternation rather than a strategic choice. Furthermore, because synchronous calls are disfavored by mate-searching females [*21*], synchrony represents a costly coordination error. Effector delay durations can vary within a species but, because they arise from fundamental physical and neurological constraints, their durations cannot be reduced to zero. Thus, synchrony is a stable feature of the selective landscape in primarily alternating choruses.

**Fig. 1.**
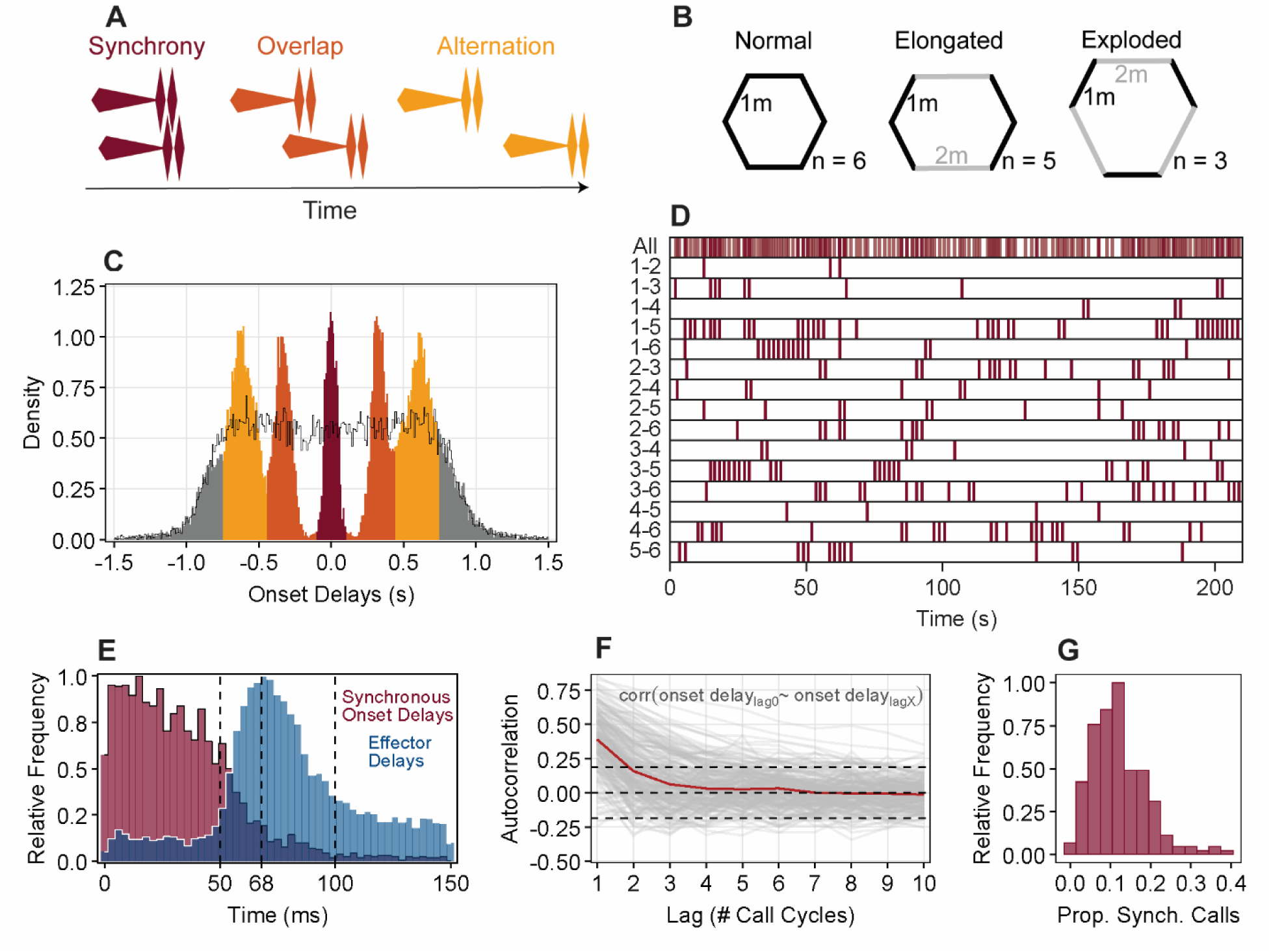
Key túngara frog chorus dynamics. **(A)** Illustrations of the discrete interaction types observed in túngara frog choruses. (**B)** Schematic diagrams of experimental chorus configurations. (**C)** Histogram (10ms bins) showing the distribution of onset delays generated by dyadic calling interactions (*n* = 24,126), and a permutation-derived null distribution (black). Bar colors match corresponding interaction types in (A). Dyadic interactions generate highly stereotyped call-timing associations. (**D)** Timelines of all synchronous calls that occurred within a chorus (top), and within each dyad (remaining rows). Dyadic synchrony clusters in time into discrete ‘synchrony runs’. (**E)** Histograms (3ms bins) of absolute onset delays between synchronous calls (burgundy; *n* = 2,864), compared to experimentally derived effector delay durations reported previously (blue; *n* = 12,117; from [*34*]). Dotted lines denote minimum (50ms) and modal (68ms) effector delays, and our current synchrony definition (100ms). Synchronous interactions bear the hallmarks of this species’ effector delay, with synchronous onset delays being shorter than the bulk of effector delay durations. (**F)** Autocorrelation plot showing correlations between dyadic onset delays in one call cycle, and onset delays at various lags. Lines for each dyad (gray; *n* = 210) and the median trend line (red) are shown. Dotted lines bound 95% confidence intervals. As can be seen, dyadic call-timing associations are highly unstable over time. (**G)** Histogram showing synchrony rate variation among dyads (*n* = 210).

Engaging in synchrony imposes attractiveness costs on males via two pathways. As stated, synchronous calls are disfavored by females [*21*; reviewed in Supplementary Table S1], meaning that males that synchronize more frequently will experience higher average costs. Furthermore, females of most frogs exhibit a perceptual bias called a ‘precedence effect’ [*29*,*30*]. As a result of this bias, when females perceive two calls closely in time, they strongly prefer the leading call over the following call (reviewed in Supplementary Table S2). In larger choruses, a key context leading one male to call within the effector delay of another, resulting in synchrony, is when multiple males’ calls are triggered by the same acoustic event [*25*,*31*]. When this happens, calls are seldom perfectly aligned in time. Thus, during synchrony one call inevitably leads the other. This makes this mutual coordination error asymmetrical, with attractiveness costs being exacerbated for the following call.

In summary, synchronous calls in alternating frogs represent coordination errors that are traceable to how a fundamental sensorimotor limitation (the effector delay) is expressed within collective social dynamics. Synchrony has important consequences for males; each synchronous call imposes baseline attractiveness costs, with the severity of these costs determined by the role occupied (leader or follower). Here, in túngara frog choruses, we mechanistically link inter-male variation in social phenotype (intrinsic calling rhythm) and sensorimotor phenotype (effector delay duration) to the frequency and severity of synchronous coordination errors. We then use experimentally derived female preference estimates to model the asymmetrical attractiveness costs that would arise from these varied error propensities. In doing so, we demonstrate the potential for phenotypically biased error outcomes to generate directional selection on sensorimotor and social phenotypes in animal collectives.

## Results

### Interaction Patterns in Túngara Frog Choruses

Male túngara frogs (*Physalaemus pustulosus*) call together in dense choruses. Their calls consist of a long ‘whine’ note followed by one or more short ‘chuck’ notes [*32*; Fig. 1A]. We recorded 84 adult males calling in 14 experimental six-male choruses. This represents an ecologically-realistic chorus size [*33*; discussion in Supplementary Text S2]. We arranged choruses in 3 shapes; ‘normal’, ‘elongated’, and ‘exploded’ hexagons (Fig. 1B).

Dyads are the most fundamental unit of collective dynamics, so we first investigated general patterns of dyadic call-timing interactions, such as the types of interactions engaged in, and the stability of dyadic call-timing associations over time. To do so, we randomly chose one male in each dyad (*n* = 15 dyads per chorus, 210 overall) to be the ‘reference male’. This designation remained consistent within dyads throughout all analyses. We then calculated ‘onset delays’ between dyad-members’ calls as the onset time of each call by the reference male minus the onset time of the closest-in-time call by the other male (*n* = 24,126 dyadic onset delays). Onset delays could be positive or negative depending on whether the reference male called first or last in a given call cycle. The distribution of dyadic onset delays showed differentiated peaks and valleys not observed in a permuted null distribution (Fig. 1C; Supplementary Text S3). Peaks correspond to three discrete interaction types that males express in larger choruses [*26*,*28*; Fig. 1A]. Males can: **1)** alternate with a rival’s call without overlapping it, **2)** overlap a rival’s call in a stereotyped way (with the onset of the following call beginning just before the chucks of the leading call), or **3)** synchronize with a rival’s call (defined here as |onset delays| < 0.1s). Autocorrelation analysis revealed that dyadic onset delays in one call cycle did not significantly predict onset delays more than a single cycle into the future (Fig. 1F). Thus, dyadic call-timing associations were highly dynamic over time with little sign of enduring rhythmic entrainment among dyad members.

Synchrony interactions (|onset delays| < 0.1s) occurred frequently and were highly stereotyped (see narrow peak around 0 in Fig. 1C; Fig. 1D). Onset delays between synchronous calls were shorter than the bulk of the effector delay distribution for this species [from *34*; Fig. 1F]. Thus, male sensorimotor systems are not capable of the rapid responses to rivals’ calls that would be needed to actively facilitate synchrony. This confirms that synchronous calls arise stochastically due to male effector delays. This contrasts with alternation and stereotyped overlap that have been shown to be adaptive interaction outcomes actively facilitated by male coordination mechanisms [*23*,*26,35*].

### Compatibility of Social Rhythms Drives Variation in Dyadic Synchrony Rates

Monte Carlo simulations revealed that dyadic synchrony rates varied non-randomly within choruses (*p* **< 0.001**; range = 0-40% of calls in synchrony; Fig. 1G; Supplementary Text S4). Like many frogs and insects, túngara frogs call rhythmically. When interactions are rhythmic, the periods of intrinsic rhythms are a key component of social phenotypes, and rhythmic compatibility can constrain interaction patterns [*36*]. Thus, we predicted that inter-male variation in intrinsic call rhythms would structure synchrony. Call periods (the intervals between onsets of successive calls by a caller) were largely stable within males (robust coefficient of variation: 11.8 ± 5 % [median ± IQR]). Thus, to summarize each male’s intrinsic call period, we calculated his median call period. Intrinsic call periods varied among males (1.78 ± 0.14s [sample mean ± SD]), and we calculated the call period difference among males in each dyad as the intrinsic call period of the reference male minus that of the other male. When doing so, intrinsic call periods for both males were recalculated using only calls produced while not calling in synchrony with one another. This removed the possibility that dyads’ tendencies to synchronize drove their call period similarity, rather than the reverse. Rates of synchronous calling were lowest for dyads with near-identical intrinsic call periods, increased at moderate-to-large differences, then decreased again somewhat for the largest differences (Bayesian-GAMM: *sds_median_* = 2.29, 95% *HDI* = [0.84, 4.3]; Fig. 2B; Supplementary Text S5).

**FIG. 2.**
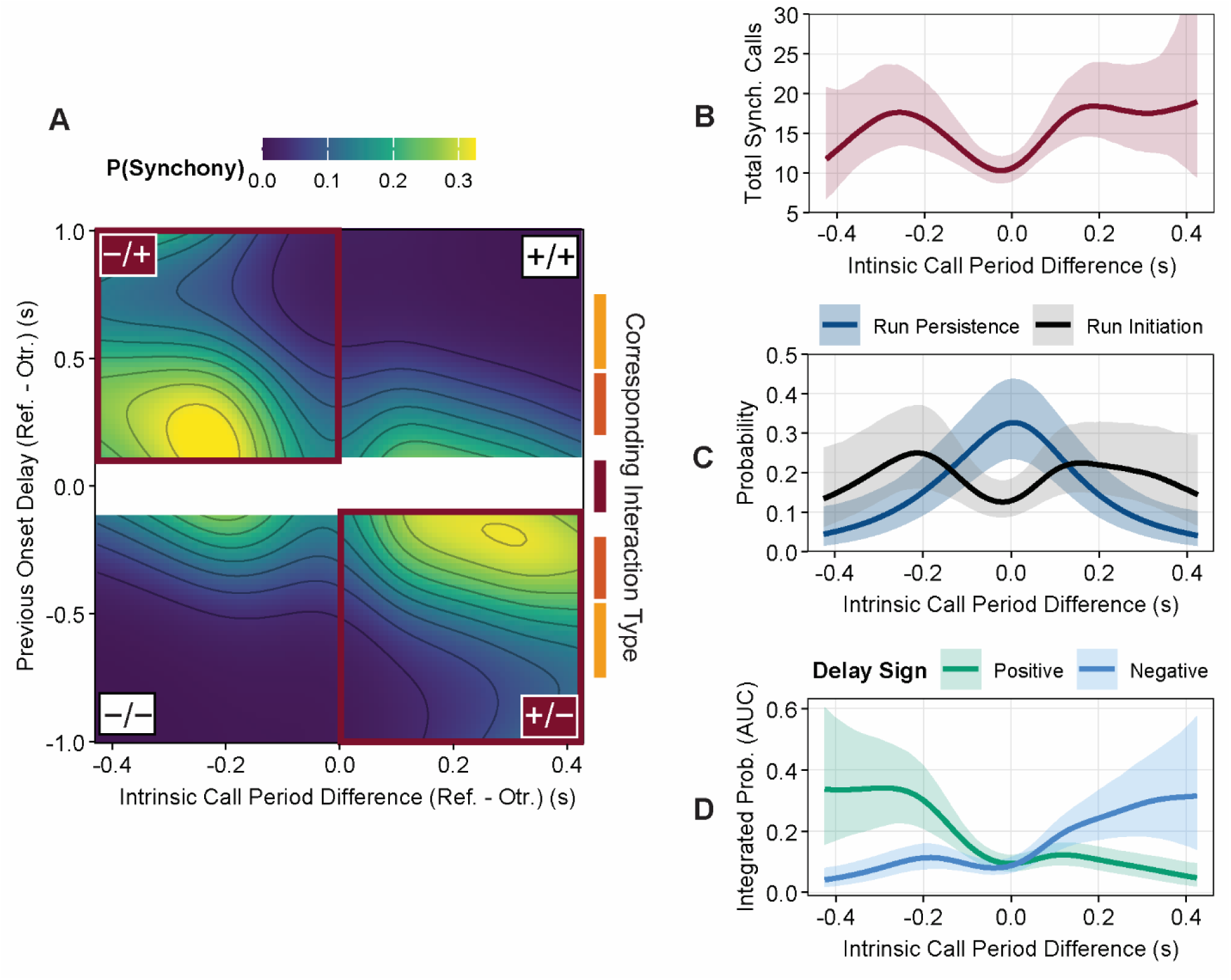
‘Temporal Offsetting’ dynamics structure dyadic synchrony patterns. (**A)** Heatmap showing the conditional effect of the linear interaction between previous onset delay and dyadic call period difference in predicting the probability that dyads calling asynchronously transitioned into synchrony (*n* = 21,382 call cycles). As predicted by our Temporal Offsetting Hypothesis, synchrony initiation probabilities were highest when dyads occupied ‘offset states’ (bounded in burgundy) in which previous onset delay and dyadic call period canceled out to some degree. Colored bars to the right illustrate the interaction types previous onset delays correspond to (see Fig. 1A). (**B)** The conditional effect of dyadic call period difference on expected dyadic synchrony counts (*n* = 210 dyads). As can be seen, dyadic call period differences structured synchrony patterns. (**C)** Conditional effects of dyadic call period difference on the probability of synchrony run initiation (*n* = 21,382 call cycles) and persistence (*n* = 2,744 call cycles), showing that dyadic call period differences exerted opposite effects on initiation and persistence probabilities. (**D)** The relationship between dyadic call period differences and average synchrony run initiation probabilities (AUC), shown separately for positive and negative previous onset delays (*n* = 210 dyads). Elevated synchrony run initiation probabilities in dyads with moderate-to-large call period differences occur exclusively when these dyads occupy ‘offset states’ (call period difference and previous onset delay carry opposite signs), demonstrating that temporal offsetting effects are the mechanism driving higher synchrony rates in dyads with larger call period differences.

### ‘Temporal Offsetting’ Explains How Rhythmic Compatibility Structures Synchrony in Time

We next investigated the mechanism by which more dissimilar intrinsic call periods increased dyadic synchrony rates. Choruses are highly dynamic, so we sought clues in how synchrony was structured in time. Dyadic synchrony was structured into ‘synchrony runs’ of back-to-back synchronous calls that varied in duration from 1-to-11 call cycles (Fig. 1D). Therefore, two factors must contribute to overall dyadic synchrony rates: **i)** how often synchrony runs were initiated, and **ii)** how long they persisted once initiated.

Since dyadic call-timing associations were highly dynamic (Fig. 1F), and variation in intrinsic call periods structured dyadic synchrony rates, we hypothesized that these factors interacted to influence the patterning of synchrony events. Males’ intrinsic calling rhythms constrain their calling behavior through time; males exhibit a refractory period after each call during which they are unable to call, followed by a gradual increase in their probability of calling again [*22,37*]. A male’s intrinsic call period thus represents the time at which his probability of calling tends to peak relative to his last call, and males varied in intrinsic call period duration. Synchrony in larger choruses often occurs when two males’ calls are triggered by the same acoustic event. This requires both males to be simultaneously occupying the latter, high-probability, phases of their respective call periods.

When dyad members’ intrinsic call periods differ in duration, aligning these latter high probability phases requires the starts of their call periods to be appropriately staggered. Consequently, associating at certain onset delays in the previous call cycle could better align these phases, increasing the probability that upcoming calls are produced synchronously. For instance, synchrony probability may be increased when the dyad-member with the longer intrinsic call period called first in the previous call cycle, providing him a head start that offsets their difference in call periods to some degree. When previous onset delays offset intrinsic call period differences in this way (i.e., when call period difference and previous onset delay carry opposite signs), we refer to a dyad as being in an ‘offset state’.

Based on this ‘temporal offsetting’ logic, we predicted that dyads calling asynchronously would have elevated probabilities of transitioning to synchrony (initiating a synchrony run) when occupying offset states. Furthermore, this effect could result in dyads with more dissimilar intrinsic call periods initiating synchrony runs more frequently, because larger call period differences might be better offset by the onset delays that arise during typical asynchronous calling in this species (∼0.3-0.6s; Fig. 1C). Regarding synchrony run persistence (remaining in synchrony from one cycle to the next), this same offsetting logic predicts that dyads with more similar intrinsic call periods will have higher probabilities of persisting in synchrony. This is because when dyad-members are already calling synchronously then, by definition, their previous onset delay was < 0.1s. Thus, alignment from this starting point is optimized when males have near-identical intrinsic call periods.

To test this, we built separate Bayesian-GAMMs to reveal how temporal offsetting influenced: **i)** the probability that dyads calling asynchronously transitioned to synchrony in the next call cycle (i.e., that they initiated a synchrony run; *n* = 21,382 call cycles), and **ii)** the probability that dyads already calling synchronously persisted in synchrony into the next call cycle (*n* = 2,744 call cycles). Models controlled for chorus shape, inter-male distances, time-varying levels of chorus activity, and attributes of dyads (Supplementary Text S6). We considered effects here robust when the Probability of Direction (*pd*) exceeded 97.5% [*38*], and considered models to differ substantially in predictive power when *Δelpd/se_diff_* > 2 [*39*].

Regarding synchrony run initiation, temporal offsetting functioned as predicted. Probabilities of transitioning from asynchronous calling to synchrony were higher for dyads occupying offset states in which intrinsic call period differences were canceled out to some degree by previous onset delays (Fig. 2A). The importance of this effect was revealed by a robust linear interaction between dyadic call period difference and previous onset delays (B-GAMM: β*_median_* = -0.62, 95% *HDI*: [-0.73, -0.52], *pd* = **100%**). Removal of this interaction diminished marginal R^2^ by 40%, and substantially decreased model predictive power (*Δelpd* = 14.9, *se_diff_* = 6, *Δelpd/se_diff_* = **2.48**). Dyadic call period differences did not influence how often dyads occupied offset states (B-GLMM: β*_median_* = -0.02, 95% *HDI* = [-0.08, 0.03], *pd* = 80%; Supplementary Text S7). However, dyads with moderate-to-large call period differences (± ∼0.2s) had higher average probabilities of transitioning to synchrony when occupying offset states (Fig. 2D; Supplementary Text S8), and so initiated synchrony runs more frequently than dyads with more similar call periods. This is likely because these call period differences are more effectively offset by the onset delays corresponding to the stereotyped overlap that predominates in 6-male choruses (onset delay = ∼0.3s; *10*; Fig. 1C; Fig. 2D).

Regarding persistence within synchrony runs once they had been initiated, dyadic call period difference had a clearly non-linear effect (B-GAMM: *sds_median_* = 1.88, 95% *HDI* = [0.87, 3.72]; Fig. 2C). Removal of this variable reduced the marginal R^2^ by 52% and resulted in a substantial reduction in predictive power (Δ*elpd* = 9.5, *se_diff_* = 4.5, *Δelpd/se_diff_* = **2.11**). As predicted, persistence probability was maximized when call period differences were near-zero (i.e., when call periods were highly similar) and diminished at higher magnitudes (Fig. 2D). Thus, once a dyad entered a synchrony run, rhythmic similarity allowed them to remain locked in synchrony for multiple call cycles.

Both the rate of synchrony run initiations and mean run duration robustly predicted overall dyadic synchrony rates (run initiation rate B-LMM: β*_median_* = 0.51, 95% *HDI* = [0.47, 0.56], *pd* = **100%**; mean run duration B-LMM: β*_median_* = 0.22, 95% *HDI* = [0.15, 0.31], *pd* = **100%**). However, most synchrony runs (70.7%) dissolved after a single synchronous call, and synchrony run initiation rate was substantially better at predicting overall synchrony rate (Δ*elpd* = 105.7, *se_diff_* = 16.7, *Δelpd/se_diff_* = **6.3**; Supplementary Text S9). Additionally, the relationship between call period difference and per-call synchrony initiation probability closely tracked that for overall synchrony rates (compare Fig. 2C to 2B). Thus, synchrony rate variation among dyads is primarily driven by varied propensities for dyads to initiate synchrony runs while calling asynchronously.

### Inter-Male Variation in Social and Sensorimotor Phenotypes Drove Synchrony Outcomes

Thus far, we have investigated the drivers of dyadic synchrony rates. However, selection on males in multi-male choruses arises from their aggregate interaction patterns with their total set of rivals. Thus, we next investigated inter-male variation in aggregate synchrony rates and role-occupation propensities (leader or follower), whether key properties of male social phenotypes (intrinsic call period) and sensorimotor phenotypes (relative effector delay duration) drove this variation, and what consequences this variation had for female preferences.

On average, 48.62 ± 11.79% [mean ± SD] of a male’s calls occurred in synchrony with at least one rival (range = 28-85%). Based on our finding that larger call period differences tended to increase dyadic synchrony rates (Fig. 2A), we predicted that males with extreme call periods would engage in synchrony at overall higher rates. The intrinsic call period (median call period) distribution for males in our sample was positively skewed (skewness = 0.76). Thus, the most extreme males were those with longer call periods. As expected, longer call periods robustly increased the overall rate at which males synchronized with rivals (B-GLMM: β*_median_* = 0.10, 95% *HDI* = [0.05, 0.15], *pd* = **100%**). This model using intrinsic call period to predict synchrony rate performed similarly to models using more chorus-specific call period deviation metrics (Supplementary Text S10).

We then investigated whether males differed systematically in their tendency to lead or follow during synchrony interactions. Synchrony often arises when multiple males have their calls triggered by the same acoustic event. In this scenario, the male with the shorter effector delay should respond more quickly and initiate his call first, and thereby lead. Thus, relative effector delay duration likely influences synchrony role. Given our observational data, we could not identify acoustic triggering events or measure effector delays. However, this logic predicts that we should see transitivity in leadership tendencies across dyads. For instance, if A tends to lead B (effector delay_A_ < effector delay_B_), and B tends to lead C (effector delay_B_ < effector delay_C_), then A should tend to lead C (effector delay_A_ < effector delay_B_ < effector delay_C_). Comparing our observed degree of transitivity (the count of non-transitive triads in our choruses) to a simulation-derived null distribution revealed that leadership propensity was indeed significantly transitive (*p* = **0.004**; Fig. 3A; Supplementary Text S11).

**FIG. 3.**
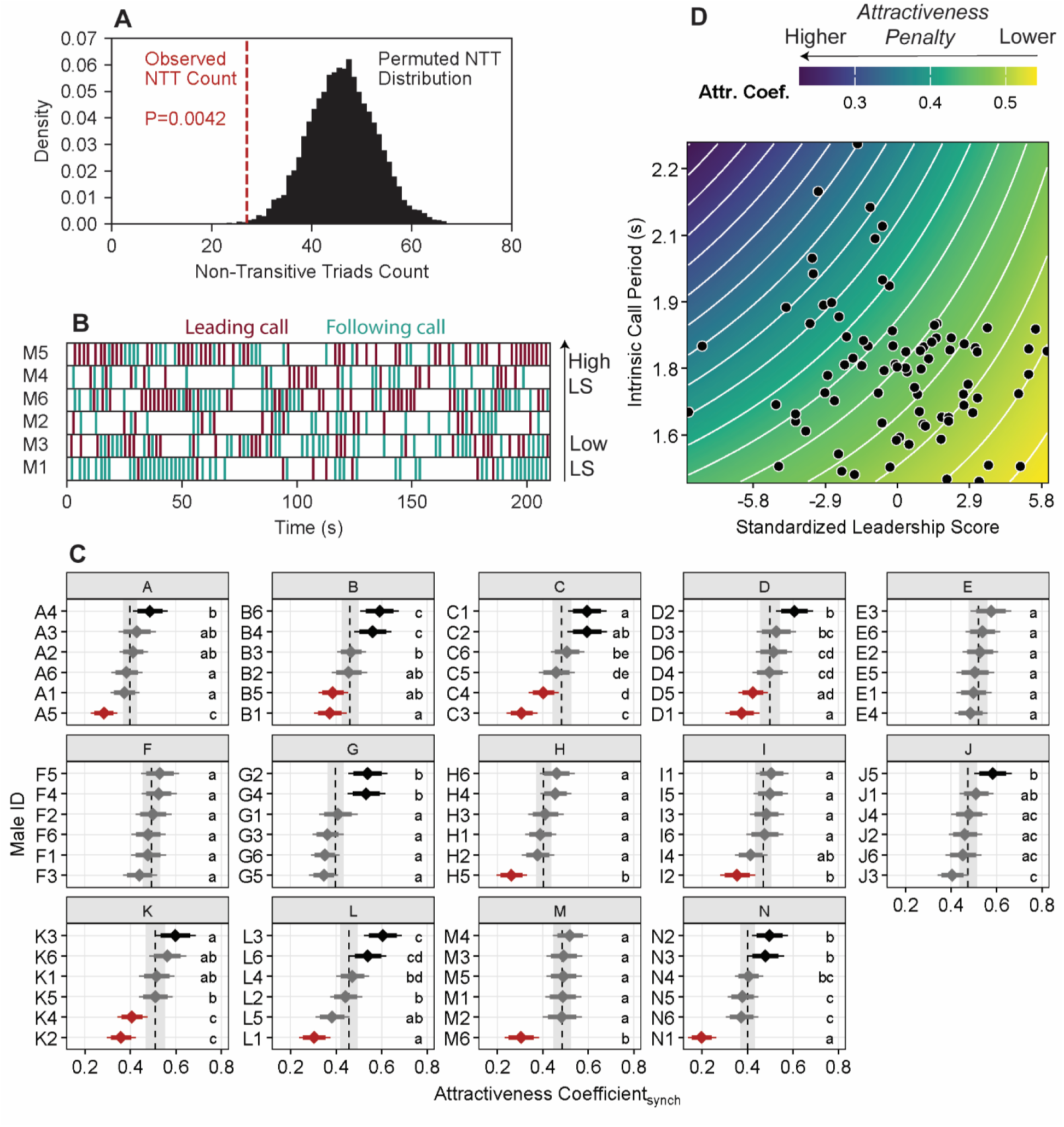
Attractiveness consequences of phenotypically biased synchrony engagement generate directional selection on sensorimotor phenotypes. **(A)** Comparing our observed leadership transitivity metric (counts of non-transitive leader/follower triads (NTT); in red) to a simulation-derived null distribution (10,000 iterations) revealed that leadership propensity is significantly transitive across dyads. This is consistent with relative effector delay durations influencing leader/follower role occupation. **(B)** Timelines showing synchronous interactions engaged in by each male in a chorus. Burgundy lines indicate occupation of the leading role, teal the following. Males are ordered vertically by leadership score (LS). **(C)** Simulated Synchrony-Derived Attractiveness Coefficient (AC_Synch_) distributions for males in each chorus (separate facets), generated via ‘Bayesian bootstrapping’ (100,000 iterations). Uncertainty is expressed by 84% and 95% CIs. Different lower-case letters denote robust inter-male differences. Black and red distributions were robustly above and below the chorus median, respectively. As can be seen, varied synchrony engagement patterns generated asymmetrical attractiveness consequences. **(D)** Conditional effects heatmap showing the predicted population-level effects of males’ intrinsic call periods and leadership scores (our proxy for effector delay duration) on AC_Synch_ score. All study males (*n* = 84) are represented as black points.

As leadership was transitive, we used methods from the dominance literature [David’s scores: *40*] to generate cardinal ‘leadership scores’ within choruses from leading/following matrices (Supplementary Text S11). We consider these a proxy for relative effector delay durations. Unsurprisingly, males with higher leadership scores had robustly lower rates of following while synchronizing (B-GLMM: β*_median_* = -0.18, 95% *HDI* = [-0.22, -0.14], *pd* = **100%**; Fig. 3B). To explore whether intrinsic call period and leadership score might be related components of a broader interactive phenotype, we investigated whether these attributes were associated. While accounting for chorus-level variation, leadership score and intrinsic call period were not robustly correlated, though exhibited a suggestive negative trend (B-MMM: r*_median_* = -0.17, 95% *HDI* = [-0.36, 0.05], *pd* = 93%; males with shorter call periods tended to have higher leadership scores (shorter effector delays); Supplementary Text S12). Neither leadership score nor intrinsic call period were robustly predicted by male body weight or length, or residual condition (52% < *pd* < 89%; all marginal R^2^ < 0.035).

### Varied Synchrony Outcomes Generate Phenotypically Biased Attractiveness Penalties

Male social phenotypes (intrinsic call period) and sensorimotor phenotypes (effector delay duration) influenced synchrony rate and role (leader/follower), outcomes known to impact female preferences [*26*,*35*]. Thus, synchrony-related female preferences may exert selection on these phenotypes. To reveal selection gradients [*41*] arising from varied synchrony engagement, we combined experimentally derived female preference strengths for non-synchronous calls over synchronous calls [*35*: P(choose non-synchronous call) = 0.83], and leading calls over following calls [*26*: P(choose leading call) = 0.94], with inter-male variation in synchrony outcomes observed in our choruses (see *Equation 1* in methods). This allowed us to calculate ‘Synchrony-Derived Attractiveness Coefficients’ (AC_Synch_) that represent multipliers applied to males’ baseline attractiveness values to account for the consequences of their expected synchrony rates and roles. However, instead of calculating a single score per male with these values, we implemented a Bayesian bootstrapping approach by sampling from distributions initialized with these values and their underlying sample sizes. This allowed us to propagate uncertainty from our behavioral observations and reported female preference values by simulating 100,000 posterior AC_Synch_ score estimates per male.

Inter-male variation in synchrony engagement generated robust variation in Synchrony-Derived Attractiveness Coefficients (AC_Synch_) in 12 of 14 choruses (Fig. 3C; Supplementary Text S13). Furthermore, our candidate phenotypes were revealed to be key proximate drivers of these varied attractiveness outcomes. As expected, intrinsic call period had a robust negative effect on AC_Synch_ scores (B-LMM: β*_median_* = -0.03, 95% *HDI* = [-0.05, -0.001], *pd* = **98%**), while leadership score had a robust positive effect (B-LMM: β*_median_* = 0.03, 95% *HDI* = [0.002, 0.06], *pd* = **98%**) (Fig. 3D; Supplementary Text S14). Their interaction was non-robust (B-LMM: β*_median_* = 0.01, 95% *HDI* = [-0.02, 0.02], *pd* = 72%). The marginal R^2^ of this model was 0.27, and this model substantially outperformed an intercept-only model in predictive power (Δ*elpd* = 73, *se_diff_* = 30.1, *Δelpd/se_diff_* = **2.4**). Thus, because longer intrinsic call periods led to higher synchrony rates, while longer effector delays increased the probability of following during synchrony, female perceptual biases against synchronous and following calls penalized these male phenotypes, likely generating selection. This demonstrates that phenotypically-biased patterns of costly coordination errors can impose directional selection on social and sensorimotor phenotypes.

## Discussion

Biological organisms possess inherent limitations which, in fast-paced social environments like animal collectives, make coordination errors inevitable. In túngara frog choruses, coordination errors in the form of inadvertent synchrony are discrete and highly stereotyped, and their costs are readily quantifiable. We leveraged this to uncover the mechanistic bases of synchrony, revealing that synchronous calls were prevalent and that the frequency and severity of these coordination errors varied by male social and sensorimotor phenotypes. Slower intrinsic call rhythms led males to engage in higher synchrony rates, while longer effector delays increased the probability that males produced costlier following calls while synchronizing. Simulations indicated that, due to female biases against synchronous and following calls, the mechanistic links between these phenotypes and synchrony outcomes should generate directional selection against longer call periods and effector delays. Thus, coordination errors here are not simply a source of random noise that imposes uniform penalties on all group members. Rather, costly error outcomes are phenotypically biased, generating winners and losers with consequences for the evolution of social strategies and their sensorimotor underpinnings.

Male frogs call to attract females to mate. Therefore, chorusing strategies have evolved to appeal to biases within female perceptual systems [*3,29*,*42*]. This includes call-timing behavior, where female biases against calls at certain temporal associations with others have selected for mechanisms that facilitate advantageous call-timing maneuvers [*3*,*29*]. Synchrony is the predominant interaction type in some chorusing insect species, and may arise as an adaptation, e.g., via beacon effects [*43*], or as a byproduct of rhythmic resetting [*3*]. Synchronous calls in alternating frogs are coordination errors that emerge when sensorimotor latencies derail call alternation strategies; *i.e.* clear byproducts. Nevertheless, they are not evolutionarily inert. The same female biases that have shaped other aspects of male calling strategy also act on synchrony outcomes, producing systematically different attractiveness consequences based on inter-male variation in synchrony rates and roles (Fig. 3D). This, in turn, should generate directional selection against the social and sensorimotor phenotypes that promoted disadvantageous synchrony outcomes, namely longer call periods and longer effector delays (Fig. 3D). In this species, expected female visitation rates [*44*], and synchronous call rates [*26*], both scale positively with chorus size. This parallel scaling means that, all else being equal, males with phenotypes that more effectively mitigate synchrony-derived costs may obtain increasingly disproportionate shares of these collective benefits. Therefore, mitigation of costly coordination errors represents no less tangible an arena for intrasexual competition than the direct competitive male-male interactions typically studied. Predation threat has also shaped chorusing [*45,46*] and other collective behaviors [*1,2*], and analogous interactions between coordination errors and perceptual biases of predators [*47*] may shape the distribution of individual risk and the evolution of collective anti-predator strategies.

Chorusing interactions are overtly rhythmic, and behavioral rhythms constrain how individuals perceive and respond to group-mate behavior [*3*,*22*,*37*]. Thus, males’ intrinsic call rhythms are a core component of their social phenotype. Despite observing a general lack of stable rhythmic entrainment within dyads (Fig. 1F), we found that rhythmic compatibility played a central role in structuring synchrony. Due to the temporal offsetting mechanism, dyads with more dissimilar call periods had higher probabilities of transitioning from calling asynchronously to calling synchronously and, consequently, engaged in synchrony at higher rates. This contrasts with synchronizing katydids, where more similar rhythms can generate more consistent synchrony in pair-wise interactions [*48*]. In the case of alternating frogs, synchronous calls represent coordination errors. Thus, our results show that rhythmic compatibility in multi-party animal collectives can not only structure target interaction patterns, but also emergent coordination errors arising from sensorimotor limitations. Behavioral rhythms [*49*], and quasi-rhythmic oscillations between behavioral states [*50*,*51*], are increasingly being shown to structure how individuals perceive and respond to group-mates’ behavior during collective locomotion in fish and insects. Our work therefore suggests that social rhythms and rhythmic compatibilities may play a deeper and more pervasive role than previously realized in shaping target and byproduct interaction outcomes in animal collectives.

Though synchrony is an inadvertent coordination error born of sensorimotor limitations, here and previously we have shown that it has tangibly influenced the evolution of male social strategies and interaction mechanisms. Emergent collective events can hone social phenotypes in two broad contexts; **1**) when the individual is participating in an event, and **2**) when the individual is external to an event. For example, in the context of escape waves in animal flocks, individuals are selected to be responsive to rival trajectories to maintain cohesion while participating in waves (context 1: [*52*]), and to glean actionable information from escape waves occurring elsewhere in the flock to, for instance, facilitate preparatory action (context 2: [*53*,*54*]). Our results here indicate that minimizing costly synchrony outcomes can exert selection on intrinsic calling rhythms and effector delay durations (context 1). Additionally, we previously showed that males attend to amplitude spikes generated by synchronizing neighbors and use them as cues when flexibly adapting their interaction strategies to current collective dynamics (context 2: [*23*]). Unlike escape waves which represent clear behavioral adaptations, synchrony is a byproduct of sensorimotor constraints; *i.e.*, an evolutionary spandrel [*sensu 55*]. Yet, synchrony as a spandrel is a consequential component of the selective landscape in choruses; both as a structured selective context to be prepared for, and as an informative social cue to attend to.

Our conclusions are based on observational data and carry three main caveats. First, we only analyzed interactions in six-male choruses, because synchrony is common in these choruses and we had a large sample size (n = 14 choruses). Though the relative frequency of the three discrete interaction types seen in túngara frogs (alternation, stereotyped overlap, synchrony) differs across chorus sizes, males’ intrinsic call periods and the structure of these interactions remain remarkably stable [*26*]. Thus, the temporal offsetting mechanism should operate similarly in any chorus large enough for synchrony to be common (>3 males) (see Supplementary Text S2 for further discussion). Second, though our simulations propagated uncertainty in both male behavior and published female preference estimates, preference estimates are from binary choice tests. Preference strengths can vary across social contexts, and so may differ in six-male choruses. However, analogous female preferences are sufficiently stable across heterogenous social contexts to have driven the evolution of precise chorusing strategies in many frogs and insects [*3,19*], and our sensitivity analysis varying the strengths of both preferences found our conclusions to be robust given that even moderate preferences persist (Supplementary Text S13). Third, when linking synchrony outcomes to sensorimotor phenotypes, we inferred the cardinal rankings of effector delay durations rather than measuring them directly. However, as would be predicted if relative effector delay durations drove leader/follower outcomes, leadership during synchrony was robustly transitive (p = 0.004). Furthermore, the David’s scores we used as proxies are designed to recover latent relative attributes from exactly this kind of observed interaction data [*40*].

## Conclusions

Coordination errors in animal collectives are often treated as noise: unstructured, undifferentiated across individuals, and therefore evolutionarily inert. However, our results overturn these assumptions. When individuals share similar sensorimotor constraints and follow similar social strategies, the same structured errors can arise repeatedly. Consequently, small differences in abilities to avoid or reduce the severity of common errors can accumulate into systematic differences in error-derived costs or risk exposure. Because errors arise synergistically from sensorimotor constraints and the social strategies within which they are expressed, selection to mitigate errors can act on social phenotypes, sensorimotor phenotypes, or both. Sensorimotor delays are universal across taxa and behaviors, and have recently been shown to shape predator-prey interactions and the evolution of predator evasion strategies [*56*]. Therefore, we suggest that coordination errors emerging from sensorimotor limitations are likely a neglected force shaping evolution within collectives more broadly. Finally, our work shows that choruses can serve as powerful ‘outsider’ systems for studying collective behavior, in which we can ask fundamental questions that are difficult to address in flocks and schools.

## Materials and Methods

### Experimental Chorus Recordings

We analyzed recordings of túngara frog males calling in fourteen 6-male experimental choruses. Choruses were recorded in 2021 and 2023 and took three spatial configurations; ‘normal’, ‘elongated’, and ‘exploded’ hexagons (see Fig. 1B for visualizations). More details regarding experimental choruses are available elsewhere [*26*]. From each chorus, we selected 3 to 3.5-minute recording segments for analysis in which all six males called stably and consistently. We extracted call onset timestamps from recordings using the Librosa [*57*] and Numpy [*58*] Python packages. See Supplementary Text S2.

### General Statistical Approach

All statistical analyses were performed in R version 4.5.1 [*59*] running in RStudio version 2025.09.01 [*60*]. For all analyses, we performed permutation/simulation tests or employed Bayesian statistical models via the ‘brms’ R package [*61*]. We fitted all Bayesian models with regularizing, weakly informative priors and standardized all continuous predictors [(x-mean(x))/SD(x)]. For statistical transparency, to minimize the risks of false positives [*62*, *63*], and to retain all covariates that might influence our main effects of interest, we did not simplify the fixed effects structure of models. Rather, we present results for full *a priori*-specified models. We report parameters as median estimates with 95% Highest Density Intervals (*HDI*). We considered effects robust when the Probability of Direction (*pd*) exceeded 97.5% [*38*], and considered models to differ substantially in predictive power when *Δelpd*/*se*_diff_ > 2 in LOOIC comparisons [*39*]. All models converged successfully (R^ < 1.01 for all parameters). Bulk and tail Effective Sample Sizes (ESS) were >500 for all fixed effects, and >300 for all group-level variance terms.

### Modeling Synchrony Run Initiation and Persistence Probabilities

We constructed separate Bayesian Generalized Additive Models (B-GAMs) to model the probability of dyadic synchrony run initiation and synchrony run persistence. Both models had a ‘Bernoulli’ outcome with a complementary log-log (cloglog) link, due to synchrony initiation and persistence events being relatively rare. We included the maximal random effects structure supported by the data [*64*]. This included a random intercept for dyad ID nested within chorus ID, as well as a multi-membership random intercept to account for the same males appearing in multiple dyads. We also initially included correlated random slopes for all time-varying dyadic covariates within dyad ID, though we removed correlations and random slopes when they were unsupported by the data (0 contained in 95% CIs). See Supplementary Text S6 for full model specifications and variable descriptions/justifications.

***Synchrony_Initiation_Model*** modeled the probability that dyads currently calling asynchronously transitioned to calling synchronously in the next call cycle (n = 21, 382 call cycles involving 210 unique dyads in 14 choruses). To test our main Temporal Offsetting Hypothesis, we included dyadic *Intrinsic Call Period Difference* and *Previous Onset Delay* as nonlinear smooths with 10 knots, and included a linear interaction between these variables. To better isolate the effect of our hypothesis of interest, we included control covariates to account for potentially salient attributes of individual males and dyads, chorus configurations, and chorus dynamics unfolding around dyad members.

***Synchrony_Persistence_Model*** modeled the probability that dyads that were already calling synchronously persisted in synchrony into the next call cycle (n = 2,744 call cycles involving 207 unique dyads in 14 choruses). This model included dyadic *Intrinsic Call Period Difference* as a nonlinear smooth with 10 knots, but excluded *Previous Onset Delay* as all dyads were already synchronous and so had previous onset delays <0.1s by definition. This model included similar control covariates as ***Synchrony_Initiation_Model***.

### Testing for Significant Transitivity of Leadership During Synchrony

To investigate whether leadership propensity during synchrony was significantly transitive across dyads, for all choruses we constructed lead/follow matrices populated with the count of synchronous interactions each male led with each other male in his chorus. We identified an overall leader for each dyad (the male that led the majority of synchronous interactions) and counted the total number of triads in which leadership was non-transitive. We then simulated a null distribution of 10,000 non-transitive triad counts that would be expected if leading/following outcomes arose purely randomly. Each iteration, we randomly assigned leadership with a probability of 0.5 for each individual dyadic synchrony interaction, while preserving the observed total number of interactions per dyad. We calculated a *p-value* as the proportion of simulated non-transitive triad counts that were less than or equal to the observed count. See Supplementary Text S11.

### Attractiveness Coefficient Calculation and Bayesian Bootstrapping

To calculate ‘Attractiveness Coefficients_Synchrony_’ (AC_Synchrony_) scores for males, we used *Equation 1* to combine male synchrony outcomes observed in our recordings (synchrony rates and leader/follower role occupation) with corresponding female preference values reported in previous phonotaxis experiments (*26*,*36*). However, we did not plug observed/reported values directly into *Equation 1*. Rather, to rigorously account for uncertainty in AC_Synchrony_ score estimates arising due to sampling effects in our current study and previous experiments, we used a Bayesian bootstrapping approach to simulate a distribution of 100,000 estimates per male.

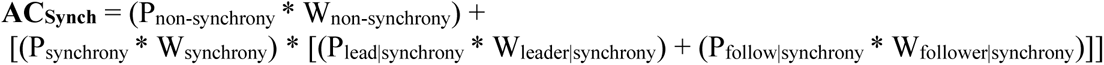

**Equation 1**: AC_Synch_ = Synchrony-Derived Attractiveness Coefficient; Psynchronous and P_non-synchronous_ = the probability that a male produced a synchronous or non-synchronous call, respectively; P_lead|synchrony_ and P_follow|synchrony_ = the probability that, during a synchronous call, a male was a leader or follower, respectively; W_non-synchronous_ = the probability weight that a female will choose a non-synchronous call over a synchronous one; W_leader_ and W_follower_ = the probability weight that, if a female chooses a synchronous, she will choose the leader or follower, respectively.

The following steps occurred each simulation iteration: ***i***) The probabilities of producing non-synchronous (P_non-synchrony_), leading synchronous (P_lead|synchrony_), and following synchronous (P_follow|synchrony_) calls for each male were drawn jointly from a multivariate Dirichlet distribution shaped by his observed calling behavior. ***ii***) Female preference weights for non-synchronous calls (W_non-synchrony_) and leading and following synchronous calls (W_leader|synchrony_, W_follow|synchrony_) were drawn from beta distributions shaped by the results and sample sizes of previous experiments (*26*,*36*). All distributions were initialized with weakly informative Laplace priors (pseudo-counts of 1). ***iii***) Values drawn from these distributions were combined via *Equation 1* to assign each male a simulated AC_Synchrony_ score. ***iv***) Pairwise differences among AC_Synchrony_ scores assigned that iteration were calculated within choruses, as were differences between each male’s assigned score and the median score of his chorus that iteration. The resulting simulated difference distributions were used to determine which differences among scores were robust. See Supplementary Text S13 for details, and sensitivity analysis.

### Calculating Sensorimotor Selection Gradients Imposed by Synchrony Outcomes

We used our Bayesian bootstrapping results to calculate selection gradients that reveal the degree to which female biases against synchronous and following calls should penalize males with sensorimotor phenotypes that generate these outcomes (slower intrinsic call periods and longer effector delays, respectively). To propagate AC_Synchrony_ score uncertainty from our simulation into our current models, we summarized each male’s score as the mean and standard deviation of his simulated score distribution. We then built a Bayesian Linear Mixed-Effects Model (B-LMM) with mean AC_Synchrony_ score as the outcome variable and with measurement error weighted by the standard deviations around these means. As fixed effects, we included intrinsic call period, leadership score (our proxy for relative effector delay duration), and their interaction. We included random intercepts and random slopes for chorus ID for all main effects. To remedy severe residual underdispersion, we fixed global residual variance (*σ*) to zero, thereby conditioning model errors entirely on the empirically derived, bootstrap-propagated measurement error. We compared predictive performance of the full model to all sub-models. Supplementary Text S14 for detailed model specifications, diagnostics, and model comparisons.

## Acknowledgements

We thank Rachel Page, Gregg Cohen, the Institute at Brown for the Environment and Society, the NOMIS Foundation, the Smithsonian Tropical Research Institute, and the University of Texas at Austin for support. We thank Guy Amichay for providing helpful comments on the manuscript, and the Peleg Lab at UC Boulder for fruitful discussions.

Research was permitted by the government of Panama (SE/A-39-2020; ARG-078-2022), and all protocols were approved by the Smithsonian Tropical Research Institute Animal Care and Use Committee (SI-21012) and the University of Texas at Austin Institutional Animal Care and Use Committee (AUP-2019-00067, AUP-2022-00012).

Google Gemini (2025/2026) was used as a sounding board while planning analysis, for generating code for data processing tasks and visualizations, and Gemini and Claude (2026) provided broad feedback on writing clarity. Google Gemini, Consensus (2025/2026) and Claude were used for literature searches. All AI-generated content was verified by the authors.

## Funding

We are grateful for funding from:

The Institute at Brown for the Environment and Society

The National Science Foundation (IOS-1914646 to MJR)

The National Science Foundation (OISE-2420214 to MJF)

The Natural Sciences and Engineering Research Council of Canada (PGS-D-567818-2022 to LCL)

The Smithsonian Tropical Research Institute

The Integrative Biology Department at UT Austin

## Author contributions

Conceptualization: LCL, MJR, MJF

Data curation: LCL

Formal analysis: LCL

Funding acquisition: LCL, MJR, MJF

Investigation: LCL

Methodology: LCL, MJR, MJF

Project administration: LCL, MJR, MJF

Resources: MJR

Software: LCL

Supervision: MJR, MJF

Validation: LCL

Visualization: LCL

Writing—original draft: LCL

Writing—review and editing: MJR, MJF

## Data accessibility

Data/code will become available upon publication.

