## supplemental for "Asymmetric Costs of Coordination Errors Generate Selection on Social and Sensorimotor Phenotypes in an Animal Collective"

**This PDF includes**:

- Supplementary Text S1-S13
- Supplementary Figures S1-S12
- Supplementary Tables S1-S19
- Protocols S1-S10

### Supplementary Text S1. Female Discrimination Against Synchronous Calls and Following Calls in Frogs

We attempted to find all published data from experimental tests of female preferences for alternating calls vs. synchronous calls (overlapping calls with call onsets within 0.1s of one another) in frogs (Table S1). Additionally, for species that produce multi-note calls, we included tests of preferences for multi-note calls with non-overlapping interdigitated notes vs. multi-note calls whose notes overlapped in a manner akin to single-note synchrony (overlapping note onsets within 0.1s). We consider both cases to be pertinent to the question of whether synchrony degrades fine-scale acoustic call structure in a way that causes the constituent calls to be discriminated against relative to non-overlapped calls. We also searched the literature for studies that experimentally tested female preferences for the leading calls vs. following calls of synchronous call pairs (Table S2).

For both tables, we only included results in which delays between the onset of synchronous calls were ≤ 0.1s (our definition of synchrony for the current study). This allowed us to focus on the effects of synchrony specifically, as distinct from more general call overlap. We also excluded data in which calls were played back in perfect synchrony (0ms between onsets of synchronous calls) as this scenario is unrealistic. When there were multiple studies for a species, or multiple relevant tests within studies, we simply include ranges of values in the relevant columns. However, we present all studies on our focal species, *Physalaemus [=Engystomops] pustulosus* as separate entries (see Supplementary Text S13 for discussion of which results we used in our attractiveness score simulations). We refer readers to the studies themselves for more detailed results.

| *Species* | *Synchronous Onset Delays Tested* | *% Females Choosing Non-synchronous Calls* | *n Per Test* | *Reference* |
|---|---|---|---|---|
| Frog Species that Actively Avoid Call Overlap | | | | |
| *Pseudacris crucifer* | 100ms | 50% | 18 | Schwartz, 1987 (1) |
| *Dryophytes versicolor* | 64ms | 79%*[N]-82%*[N] | 14-17 | Schwartz, 1987 (1) |
| *Dendropsophus microcephala* | 12ms | 63%*[N]-93%*[N] | 14-38 | Schwartz, 1987 (1); Schwartz, 1993 (2) |
| *Dryophytes avivoca* | 20-60ms | 22%*[S]-82%*[N] | 16-19 | Martínez-Rivera & Gerhardt, 2008 (3) |
| *Physalaemus pustulosus* | 79ms | 82.5%*[N] | 40 | Legett et al., 2019 (4) |
| *Physalaemus pustulosus* | 50-100ms | 35-65% | 20 | Schwartz & Rand, 1991 (5) |
| Frog Species that Actively Overlap/Synchronize their Calls | | | | |
| *Centrolenella granulosa* | 100ms | 55%-85%*[N] | 11-13 | Ibanez, 1993 (6) |

**Table S1**. Preferences for Non-Synchronous Calls in Frogs. *[N] denotes significant preference for non-synchronous calls over synchronous calls; *[S] denotes significant preference for synchronous calls over non-synchronous calls. Green shading indicates overall support for preference for non-synchronous calls, orange indicates overall lack of support.

| *Species* | *Onset Delays Tested* | *% Females Choosing Leading Call* | *n Per Test* | *Reference* |
|---|---|---|---|---|
| Frog Species that Actively Avoid Call Overlap | | | | |
| *Hyperolius marmoratus* | 0.5-70ms | 81%$*^{L}$-100%$*^{L}$ | 14-16 | Grafe, 1996 (7); Dyson & Passmore, 1988 (8,9) |
| *Dryophytes versicolor* | 2-18ms | 100%$*^{L}$ | 12 | Marshall & Gerhardt, 2010 (10) |
| *Dryophytes cinereus* | 25-100ms | 85%$*^{L}$-100%$*^{L}$ | 20-42 | Reichert et al., 2016 (11); Höbel and Gerhardt, 2007 (12) |
| *Hyla chrysoscelis* | 0.5-5ms | 85%$*^{L}$-100%$*^{L}$ | 20 | Krishnan & Reichert, 2026 (13) |
| *Physalaemus pustulosus* | 50-79ms | 90$*^{L}$-94%$*^{L}$ | 35-40 | Legett *et al.* , 2020 (14); Larter & Ryan, 2024 (15) |
| *Physalaemus pustulosus* | ≥10ms | Not stated$*^{L}$ | Not stated | cited in: Greenfield and Rand, 2000 (16) |
| *Physalaemus pustulosus* | 50-100ms | 50-54% | 20 | Schwartz & Rand, 1991 (5) |
| Frog Species that Actively Overlap/Synchronize their Calls | | | | |
| *Centrolenella granulosa* | 100ms | 57% | 7 | Ibanez, 1993 (6) |
| *Kassina fusca* | 18.5-92.5ms | 5%$*^{F}$-37% | 13-16 | Grafe, 1999 (17) |
| *Smilisca sila* | 79ms | 65% | 23 | Legett *et al.* , 2020 (14) |

**Table S2**. Precedence Effects in Frogs. $*^{L}$ denotes significant preference for leading calls of synchronous pairs; $*^{F}$ denotes significant preference for following calls. Green shading indicates overall support for preference for a precedence effect, orange indicates overall lack of support.

#### *Trends in the Literature*

Table S1 shows that, in the majority of alternating frogs tested, females show a significant preference for alternating calls over synchronous calls. This is likely because important elements of the fine temporal structure of calls can be degraded when calls synchronize and overlap (*18,19*). Table S2 shows that, when calls are synchronous, females of most alternating frogs show a strong preference for leading calls over following calls. The demonstrates that precedence effects are a widespread sensory bias shaping female mate choice decisions (*20*).

### Supplementary Text S2. Chorus Recording Methods

#### *Experimental Chorus Recordings and Timestamp Extraction*

All experimental choruses consisted of males arranged in hexagonal configurations. 'Normal' and 'elongated' choruses were recorded in Fall of 2021, while all 'exploded' choruses were recorded in Fall of 2023 (see Fig. 1B, in-text, for visualizations of these chorus configurations). All recordings took place in a darkened room (more details in: *15*) with ambient temperatures ranging from 25C to 27.3C. Each male within each 6-male experimental choruses called within a separate acoustically transparent enclosure that was ~11.4cm in diameter. These enclosures were centered on the vertices of these hexagonal configurations. Enclosures contained water on which males could float to call. Each male was recorded onto a separate audio track of a

Zoom F6 recorder via a separate SYNCO lav-S6 tie clip microphone attached to his enclosure. For this study, we selected 3 to 3.5-minute segments from chorus recordings in which all six males called stably and consistently, and we used call timestamps extracted from these segments for analysis. To extract the timestamps of call onsets, we used functions from the Librosa (*21*) and Numpy (*22*) Python packages to extract the peak amplitude envelope of each caller's audio track. From this, we located amplitude peaks corresponding to calls and worked backwards from these peaks to obtain precise timestamps for call onsets. All onset timestamps were visually verified during processing.

***Justification for Focusing on 6-male Choruses for this Study***

Though in 2021 we recorded choruses ranging in size from 2-6 males (*15*), for this current study we only focused on six-male choruses. We made this choice for several reasons. For one, this study focused on synchronous interactions, and synchrony rates increase with chorus size. Six was the largest chorus size we recorded and so had the highest synchrony rates, giving us the best sample set for revealing the dynamics that lead to synchrony. Furthermore, we desired a standardized social environment in which we could detect the potentially subtle mechanistic effects we were interested in. Of all chorus sizes we possessed data for, we had the largest sample size for 6-male choruses (n = 14 six-male choruses vs. n = 5-8 for other chorus sizes). This was because the 'extended' and 'exploded' hexagonal choruses were only recorded as 6-male choruses. Finally, we decided to focus solely on 6-male choruses, rather than pooling data from all choruses regardless of size, because our GAMM models for detecting the mechanisms driving synchrony were highly complex and computationally demanding (see Supplementary Text S6). Therefore, accounting for the varied dynamics across varied chorus sizes in a statistically-appropriate manner would have been analytically intractable. This would have required complex model-wide interactions with chorus size or, at least, many pairwise interactions. However, we believe that the dynamics we reveal here will be operating in any large chorus in which synchrony occurs regularly (>3 males), which we explain below.

Though túngara frogs call in varied social environments and social dynamics vary in certain ways across these environments, we believe six-male choruses provide a robust and representative model for evaluating the broader mechanistic basis of synchronous interactions. This is because key elements of the Temporal Offsetting phenomenon we demonstrate here do not vary in form with chorus size. For one, call periods are stable within males across all tested chorus sizes (2-6 male choruses: *14*). Furthermore, though interaction modes (alternation, overlap, synchrony) vary in prevalence across chorus sizes, the temporal form that these interactions take, and thus the onset delays that arise from them, remain remarkably stable across chorus sizes (*15*). Thus, though synchrony rates vary across chorus sizes, the synchrony-promoting mechanisms of rhythmic compatibility and temporal offsetting revealed here should be structuring the synchrony that does occur. As a final note on the generalizability of our results, we also point out that in the GAMM models below in which we modeled the temporal offsetting effect, there was no effect of male spatial geometry on the probability of dyadic synchrony run initiation or persistence. Neither the shape of the chorus (hexagonal, elongated, exploded), nor whether dyad members were nearest neighbors or not, nor their interaction, had a robust effect in either the synchrony run initiation or persistence model (Supplementary Text S6). Conversely, predictors describing our temporal offsetting hypothesis were highly robust. Thus, the robust temporal offsetting effects shown in these models arise in spite of variation in spatial configurations, suggesting these effects are generalizable across different chorus organizations.

### Supplementary Text S3. Generating a Permutation-Derived Null Distribution of Dyadic Onset Delays

To ensure that the clear patterns arising in this species' onset delay distribution (Fig. 1C, in-text) could not arise by chance, we also generated a null distribution of dyadic onset delays with which to compare our observed distribution. We did so by permuting the call periods of each male in each chorus and building new call sequences from these permuted call period sequences. This then gave us a permuted series of call timestamps for each male that preserved his call period distribution and therefore his typical calling rhythms, but divorced the precise timing of his calls from the timing of the calls of his chorus-mates (*23*). With this permuted dataset, we calculated dyadic onset delays between calls in exactly the same way described in-text: 'we randomly chose one male in each dyad (n = 210) to be the 'reference male' and calculated 'onset delays' between dyad-members' calls as the onset time of each call by the reference male minus the onset time of the closest-in-time call by the other male. Onset delays could be positive or negative depending on whether the reference male called first or last in a given call cycle.' We plot this permuted null distribution in black in Fig. 1C, in-text. As can be seen, observed onset displays deviate markedly from the null expectation.

- *Code for function to permute call periods found in Python Jupyter notebook 'Dyadic_DTSM_Data_Wrangling_FINALIZED.ipynb'.*
- *Code for plotting observed and permuted onset delay distribution found in RMarkdown file 'SYNCH_FIGS_FINALIZED.rmd'.*

### Supplementary Text S4. Testing for Robust Variation in Dyadic Synchrony Rates Within Choruses

To test whether variation among dyads in their synchrony rates differed from random expectations, we performed Monte Carlo simulation tests. For each chorus, we calculated the chorus-specific expected probability of synchrony as the overall rate of synchrony in that chorus (count of synchronous dyadic call cycles in the chorus divided by the sum of all synchronous and asynchronous dyadic call cycles). We then calculated the observed sum of squared deviations (SSD) by comparing each dyad's observed synchronous call cycle count to its expected count (the total number of call cycles performed by given dyad multiplied by the baseline synchrony probability of their chorus). We then ran 1,000 simulations in which each dyad's synchrony count was generated via a binomial distribution (n = observed number of call cycles, p = chorus-specific expected rate). For each iteration, we calculated the SSD across all dyads. The *p-value* was calculated as the proportion of simulated SSDs that were greater than or equal to our observed SSD. This revealed that the observed variation in dyadic synchrony rates was significantly higher than expected under the null ($p < 0.001$; Fig. S1, below).

- *Code for performing and visualizing this simulation found in RMarkdown file 'SYNCH_FIGS_FINALIZED.rmd'.*

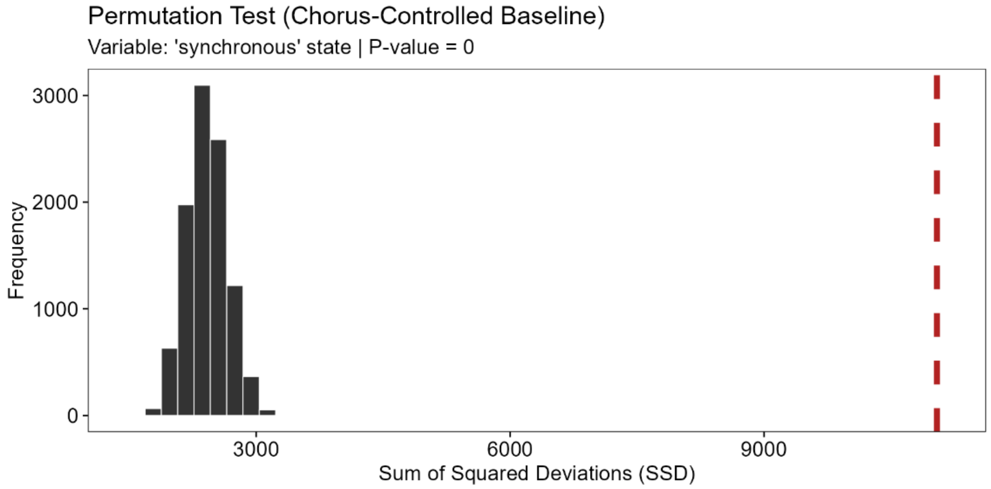


**FIG. S1**. Results of Monte Carlo simulation test. Observed SSD (red dashed line) was higher than all simulated null SSDs (black histogram), indicating that dyads exhibit meaningful variation in synchrony rates.

**Supplementary Text S5. Investigating the Relationship Between Intrinsic Call Period Differences and Overall Dyadic Synchrony Rates**

We investigated whether synchrony rates were structured by differences among males in their intrinsic call periods. In Fig. S2 below we show the distribution of intrinsic call period (median call periods) among males and the distribution of absolute intrinsic call period differences within dyads. We took the absolute value here as the sign of this difference was arbitrary and purely based on which male was the reference male and which was the other male (see Supplementary Text S6).

- *Code for plotting these histograms found in RMarkdown file 'SYNCH_FIGS_FINALIZED.rmd'.*

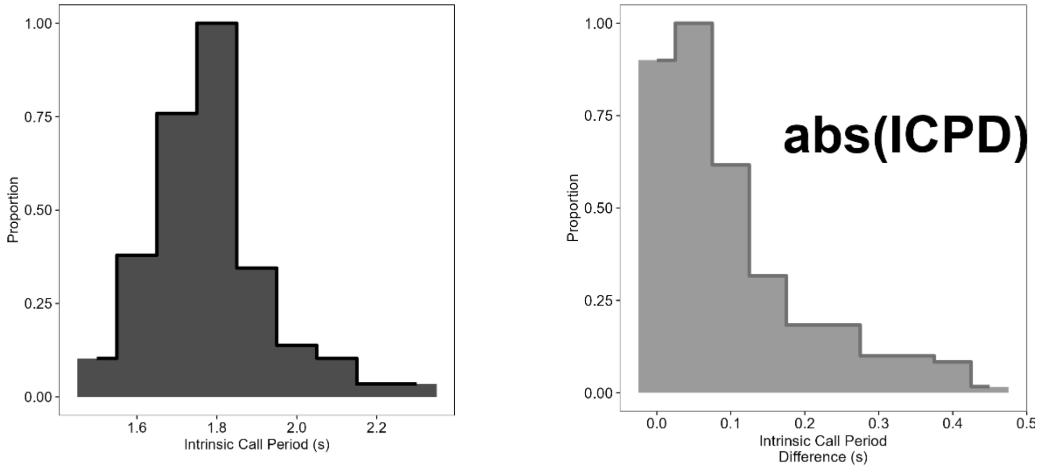


**FIG. S2.** *Left*: distribution of intrinsic call periods (ICP) among males (median call periods; n = 84). Right: distribution of absolute Intrinsic Call Period Differences (ICPD) of dyads (n = 210 across 14 choruses). ICPD could be positive or negative, but we show the absolute value here.

***Modeling the Effect of Intrinsic Call Period Difference on Dyadic Synchrony Rates***

To see how differences among dyad members in intrinsic call periods structured their synchrony rates, we constructed a negative binomial Bayesian GAMM in 'brms' (*24*) (syntax and results below; Protocol S1; Table S3). We included the count of the total number of synchronous calls dyads engaged in as the response variable, and intrinsic call period differences between dyad members as the predictor variables; we modeled this relationship with a smooth with 10 knots as intrinsic call period differences could be positive or negative (see Supplemental Text S6 for calculation of this variable) and we expected high magnitude positive and negative differences to have similar effects that differed from those near zero. We also included an offset (log(Synchrony_Opportunities)), with Synchrony_Opportunities being the number of calls produced by the dyad member that called least during the observation period, as this represents the number of opportunities the dyad had to potentially synchronize. In Fig. 2B, in-text, we show predictions for dyads with the median Synchrony_Opportunities seen in our dataset.

- *Code for this model, model checks, and model results in RMarkdown file 'DYADIC_SYNCH_OUTCOMES_FINALIZED.rmd'.*

```
frog_formula_nb <- bf(
  # Main Model
  Total_Synchronous_Calls ~ s(Dyadic_Int_Call_Period_Diff_std, k = 10) + offset(log(Synchrony_Opportunities)) +
(1 | Chorus_Name),

  # Dispersion Model (shape) to handle complex variance
  shape ~ Dyadic_Int_Call_Period_Diff_std + (1 | Chorus_Name)
)

# Run the standard negative binomial model
model_count_nb <- brm(
  formula = frog_formula_nb,
  data = data,
  family = negbinomial(link = "log"),
  chains = 4,
  cores = 4,
  threads = threading(2),
  iter = 2000,
  warmup = 1000,
  backend = "cmdstanr",
  control = list(
    adapt_delta = 0.999,
    max_treedepth = 20
  )
)
```

**Protocol S1**. Brms syntax for our model investigating the effect of dyadic call period difference on synchrony rate.

#### *Model Results*

| Parameter | Median | 95% HDI Lower | 95% HDI Upper | pd | R-hat | ESS |
|---|---|---|---|---|---|---|
| ***Population-Level (Fixed) Effects*** | | | | | | |
| b_Intercept | -2.142 | -2.255 | -2.021 | 1.000 | 1.002 | 2,241 |
| b_shape_Intercept | 1.643 | 1.185 | 2.144 | 1.000 | 1.001 | 2,097 |
| b_shape_Dyadic_Int_Call_Period_Diff_std | -0.062 | -0.403 | 0.264 | 0.649 | 1.001 | 4,997 |
| bs_sDyadic_Int_Call_Period_Diff_std_1 | 2.612 | -2.688 | 8.716 | 0.842 | 1.001 | 2,294 |
| ***Group-Level (Random) Effects (SD)*** | | | | | | |
| sd_Chorus_Name__Intercept | 0.144 | 0.018 | 0.284 | 1.000 | 1.000 | 855 |
| sd_Chorus_Name__shape_Intercept | 0.564 | 0.002 | 1.110 | 1.000 | 1.002 | 848 |
| ***Smooth Term Variances (Wiggliness)*** | | | | | | |
| sds_sDyadic_Int_Call_Period_Diff_std_1 | 2.287 | 0.837 | 4.309 | 1.000 | 1.001 | 1,487 |

**Table S3**. Medians and upper and lower 95% *HDI*s of posterior distributions of fixed effects, random effects, and smooth terms. Also shown are Probability of Direction (*pd*), R-hat, and effective sample size.

Our model revealed a robust, non-linear relationship between intrinsic call period difference and synchronous calling behavior (sds = 2.29, 95% *HDI* = [0.84, 4.31]). Visually (Fig. 2B, in-text), it can be seen that dyadic synchrony rates were lowest for dyads with near-identical call periods (intrinsic call period = ~0) and increase at moderate-to-large intrinsic call period differences (~0.2s), then decreased somewhat at the largest differences.

### Supplementary Text S6. Modeling Approach for Uncovering Mechanisms Promoting Synchrony

#### *Notes on Our General Statistical Approach in this Study*

All analyses were performed in R version 4.5.1 running in RStudio version 2025.09.01. We employed a Bayesian framework for all statistical modelling. We fitted all Bayesian models using the 'brms' (*24*) R package and used regularizing, weakly informative priors. For all models, we standardized continuous predictors [(x-mean(x))/SD(x)] prior to modeling. For statistical transparency, to minimize the risks of false positives (*25*, *26*), and to retain all covariates that might confound our main effects of interest regardless of the magnitude of their effects, we did not simplify the fixed effects structure of models. Rather, we present results for full *a priori*-specified models. We report parameters as median estimates with 95% Highest Density Intervals (*HDI*). We considered effects robust when the Probability of Direction (*pd*) exceeded 97.5% (*27*), and considered models to differ substantially in predictive power when $\Delta elpd/se_{\mathrm{diff}} > 2$ (*28*). All models converged successfully ($\hat{R} < 1.01$ for all parameters). Bulk and tail Effective Sample Sizes (ESS) exceeded 500 for all fixed effects, and were >300 for all group-level variance terms.

#### *Bayesian GAMM Framework*

Dyadic synchronous interactions were clearly structured in time, with synchrony occurring in continuous synchrony 'runs' of 1 or more back-to-back synchronous calls (range = 1-11; see Fig. 1D, in-text). We predicted certain variables would influence both the probability that dyads initiated synchrony runs (transitioned from calling asynchronously to synchronously) and the probability that synchrony runs would persist once initiated (dyads calling synchronously would continue in synchrony into the next call cycle). Furthermore, we predicted that certain variables would influence each of these probabilities in different ways due to the logic of our Temporal Offsetting Hypothesis (explained in-text). Thus, we constructed separate models to model the probability of synchrony run initiation and persistence, which we refer to as *Synchrony_Initiation_Model* and *Synchrony_Persistence_Model*, respectively.

We constructed Bayesian Generalized Additive Models. These models had a 'Bernoulli' outcome with a complementary log-log (cloglog) link due to synchrony initiation and persistence events being relatively rare overall. We utilized regularizing, weakly informative priors in both models to prevent overfitting on cloglog scale. We also included the maximal random effects structure supported by the data (*29*). This included a random intercept for dyad ID nested within chorus ID to account for unexplained dyad-level and chorus-level variation, as well as a multi-membership random intercept which accounted for unexplained variation among individual males in their interaction propensities when members of different dyads. We also initially included correlated random slopes for all time-varying dyadic covariates within dyad ID, though we removed correlations and random slopes if they were unsupported by the data (0 contained in 95% CIs).

### ***SYNCHRONY INITIATION MODEL***

This model investigated the factors influencing the probability that dyads calling asynchronously transitioned into synchrony in the next call cycle. Thus, each row of the dataframe represented a dyad *whose most recent calls had been produced asynchronously (not in synchrony)*, and the outcome variable was a binary response denoting whether their next calls occurred in synchrony (1; synchrony was initiated) or not (0). For this model, n = 21, 382 call cycles involving 210 unique dyads, in 14 choruses. We describe the variables included in this model, present the model syntax, and present model results, below.

- *Code for this model, model checks, and model results in RMarkdown file 'GAMM_MODELS_FINALIZED.rmd.*
- *Code for generating visualizations of model results in RMarkdown file 'GAMM_VIZ_FINALIZED.rmd.*

#### ***Fixed Effects: Synchrony_Initiation_Model***

We generated dyadic covariates, described below, for all possible dyad combinations in each chorus. We randomly selected one male from each dyad to be the '*Reference Male'* (*RM*) and designated the remaining male the '*Other Male*' (*OM*). We calculated all dyadic covariates from the 'point of view' of the *Reference Male* (indicated by '*RM*' or '*OM*' subscript). The designations of *RM* and *OM* within dyads were fixed and never changed across contexts in this study.

##### *Dyadic Fixed Effects Representing the Temporal Offsetting Hypothesis*

Key dyadic fixed effects for testing the Temporal Offsetting Hypothesis were:

- *Intrinsic Call Period Difference:* The difference in the period of dyad members' intrinsic calling rhythms. To represent males' intrinsic call periods, we calculated their median call period throughout the observation period. When doing so, intrinsic call periods for both males in each dyad were recalculated using only calls produced while not calling in synchrony with one another. This removed the possibility that synchrony led to call period similarity rather than the reverse. Intrinsic call period difference was then calculated as: *Intrinsic Call Period*$_{RM}$ *- Intrinsic Call Period*$_{OM}$.

- *Previous Onset Delay:* This denoted how dyad members' calls were temporally associated in the call cycle that immediately preceded the call of interest. Again, these were calculated from the POV of the reference male, and so were calculated as the onset time of the immediately preceding call by the reference male minus the onset time of the closest-in-time call by the other male: *Onset Time of Previous Call*$_{RM}$ *- Onset Time of Closest-In-Time Previous Call*$_{OM}$. Dyadic call-timing associations changed constantly over time (Fig. 1F, in-text), and so this was calculated anew each call cycle as a time-varying variable.
- The *Temporal Offsetting Hypothesis* predicted that *Intrinsic Call Period Difference* and *Previous Onset Delay* would interact, with synchrony initiation probabilities being highest when intrinsic call period differences were offset to some degree by previous onset delays; i.e., when these quantities had opposite signs. Thus, the true test of this hypothesis was achieved via including a linear interaction between these variables (see model syntax below).

*Dyadic Control Variables*

Certain additional variables needed to be controlled for as they might also influence a dyad's propensity to synchronize. For instance, in túngara frogs, calling behavior is influenced somewhat by certain male attributes (*30*), and males that are nearer to one another in the chorus will experience a more similar view of the fluctuating acoustic scene at the chorus. Thus, similarity in key attributes, or spatial proximity, could lead males to make more similar call-timing decisions. This, in turn, might increase the probability that dyad members select the same moment in time to call, increasing the probability of synchrony. So we controlled for these variables by including them as covariates:

- *Mean Call Period Variability*: Call period variability could influence how reliably our Temporal Offsetting mechanism could function, as increased variability adds noise to the system. Thus, it had to be controlled for. Call period variability for each male was calculated using a variant of the Root Mean Square of Successive Differences metric used to measure heart rate variability (*31*). However, we used the median of successive differences, rather than the mean, to increase robustness to outlier call periods. Thus, variability for each male was calculated as the square root of the median squared differences of successive call periods. Call period variability for males in each dyad was recalculated using only calls produced while not calling in synchrony with one another. This removed the possibility that dyadic synchrony influenced call period variability rather than the reverse. Mean Call Period Variability for the dyad was then calculated as: *mean(Call Period Variability*$_{RM}$*, Call Period Variability*$_{OM}$*)*.
- *Weight Difference*: Calculated as: *abs(body weight*$_{RM}$ *– body weight*$_{OM}$*)*.
- *Chorus Shape*: A categorical variable denoting the shape of the chorus configuration: 'normal', 'elongate', or 'exploded' (see Fig. 1B, in-text).
- *Nearest Neighbors*: A binary variable denoting whether males were nearest neighbors ('Nearest'; when males were within 1m of one another), or not. As the relative importance of nearest neighbors likely differs in our different chorus configurations due to their different inter-male distance distributions, we also included an interaction between *Chorus Shape* and *Nearest Neighbors*.

- *Leadership Score Difference*: Leadership scores are described in-text and we consider them a proxy for the relative durations of males' effector delays. As synchrony arises as a result of male effector delays, we thought similarity in these scores could influence synchrony. We calculated a dyad's Leadership Score Difference as: *abs(leadership score*$_{RM}$ *– leadership score*$_{OM}$*).*

*Control Variables Pertaining to Broader Chorusing Dynamics*

Frogs, including túngara frogs, are responsive to the dynamics of their local social environments and this can influence their interaction tendencies(*30*, *32*). Chorus dynamics are heterogeneous through time, and we thus needed to control for these dynamics when analyzing each interaction. Thus, we included time-varying variables denoting current chorus dynamics as control variables:

- *Instantaneous Chorus Size*: A measure of current activity levels in the broader chorus. Calculated as the count of the total number of calls produced by all other males at the chorus (outside of the focal dyad) during the call period of the reference male leading up to the current interaction (max value = 4 in our 6-male choruses).
- *Effective Chorus Size*: A measure of the duty cycle of chorusing activity in the broader chorus. The same as *Instantaneous Chorus Size*, but with synchronous calls by rivals counted as one single combined call (max value = 4, always ≤ *Instantaneous Chorus Size*).
- We also included an interaction between these variables which functioned as a proxy for whether other males in the chorus were synchronizing. This is because when *Effective Chorus Size* is less than *Instantaneous Chorus Size*, this indicates males elsewhere in the chorus must be synchronizing.

*Temporal Control Variables*

- *Asynchrony Run Duration*: To control for the possibility that the probability that a dyad synchronizes changes in a systematic way through time as they call asynchronously, we included *Asynchrony Run Duration* as a time-varying covariate. This variable tracked the current number of call cycles for which the dyad had been continuously calling asynchronously; i.e., the duration in call cycles of the current run of asynchronous calling, updated each call cycle.
- *Run Start Time Known*: As the first asynchronous calling run began before the start of our observation period, we did not know its true duration. Thus, we included *Run Start Time Known* as a binary variable denoting whether an asynchrony run was the initial run or not. We also included an interaction between this variable and *Asynchrony Run Duration*, to allow us to only present model results regarding the effects of duration for runs with known durations.
- *Intrinsic Call Period of the Reference Male*: As *Asynchrony Run Duration* is measured in the counts of call cycles, the comparability of the effect of time on synchrony probability across dyads could be compromised if dyads differ greatly in the intrinsic call period of their reference male. Thus, we also controlled for the *Intrinsic Call Period of the Reference Male*, to make run durations comparable across dyads.

***Modeling Approach: Synchrony_Initiation_Model***

Because we expected the effects of *Intrinsic Call Period Difference* and *Previous Onset Delay* to be non-linear, we included these variables as nonlinear smooths with 10 knots. As both variables were calculated from the 'point of view' of the reference male ($value_{reference\_male}$ – $value_{other\_male}$), they could take both positive or negative values depending on the relative magnitude of the value of the reference male and other male. Our temporal offsetting hypothesis predicted that certain combinations of *Intrinsic Call Period Difference* and *Previous Onset Delay* would cancel each other out to promote synchrony onset. Thus, we also included a linear interaction between these variables. As control covariates, we included linear terms for *Mean Call Period Variability, Weight Difference*, and *Leadership Score Difference*. To control for spatial influences, we included whether males were *Nearest Neighbors,* and *Chorus Shape,* as unordered factors, and their interaction. To capture general chorus dynamics occurring around dyad members, we included *Instantaneous Chorus Size*, *Effective Chorus Size*, and their interaction, as proxies for overall duty cycles and degrees of synchrony elsewhere in the chorus (if *Effective Chorus Size* < *Instantaneous Chorus Size* this indicates synchrony elsewhere in the chorus). Finally, to control for temporal effects we included the current duration of the current asynchronous call run (*Asynchrony Run Duration* in interaction with *Run Start Time Known*) and controlled for the *Intrinsic Call Period of the Reference Male* to make asynchrony run durations comparable across different dyads. Model syntax is shown in Protocol S2, results in Table S4.

```
#Regularizing, weakly informative, priors
custom_priors_final <- c(
    # Fixed Effect Prior
        set_prior("normal(0, 0.5)", class = "b"),
    # Intercept Prior
        set_prior("normal(0, 3)", class = "Intercept"),
    # Random Effect SDs Prior
        set_prior("exponential(1)", class = "sd"),
    # Smooth Term SDs Prior
        set_prior("cauchy(0, 0.5)", class = "sds"))

#Model
Synchrony_Initiation_Model <- brm(synchrony_onset_event ~
    #Test of Temporal Offsetting Hypothesis
            s(Intrinsic_Call_Period_Difference_std, k = 10)+
            s(Previous_Onset_Delay_std, k = 10)+
            Intrinsic_Call_Period_Difference_std : Previous_Onset_Delay_std +        #interaction for 'temporal offsetting'
    #Dyadic Control Variables
            Mean_Call_Period_Variability_std +
            Leadership_Score_Difference_std +
            Weight_Difference_std +
            Nearest_Neighbors * Chorus_Shape+
    #Broader Chorus Dynamics Control Variables
           Effective_Chorus_Size_std * Instantaneous_Chorus_Size_std +
    #Temporal Control Variables
            Asynchrony_Run_Duration_std * Run_Start_Time_Known +
            Intrinsic_Call_Period_of_Reference_Male_std +
    #random effects
           (1 | Chorus_Name/Dyad_ID) +                                  #dyad_ID nested in chorus_ID
           (0 + Previous_Onset_Delay_std +
             Effective_Chorus_Size_std +
             Instantaneous_Chorus_Size_std +
             Asynchrony_Run_Duration_std || Chorus_Name : Dyad_ID) + #uncorrelated random slopes
           (1 | mm(Reference_Male_ID, Other_Male_ID)),       #mulitmembership term for individual contributions
    #Run settings
  data = data,
  family = bernoulli(link = "cloglog"),
  prior = custom_priors_final,
  init=0.1,                                      #restrict initial values to stabilize cloglog link initialization
  chains = 4,
  cores = 4,
  threads = threading(2),
  iter = 2700,
  warmup = 1500,
  refresh = 5,
  backend = "cmdstanr",
  control = list(
    adapt_delta = 0.999,                         # small step size to eliminate divergent transitions
    max_treedepth = 15                           # Deeper tree exploration
)
```

**Protocol S2**. Brms syntax for Synchrony_Initiation_Model.

#### *Model Results: Synchrony_Initiation_Model*

| Parameter | Median | 95% HDI Lower | 95% HDI Upper | pd | R-hat | ESS |
|---|---|---|---|---|---|---|
| ***Population-Level (Fixed) Effects*** | | | | | | |
| **b_Intercept** | -3.461 | -3.803 | -3.125 | 1.000 | 1.001 | 2,968 |
| b_Dyadic_Call_Period_Var_std | 0.046 | -0.077 | 0.174 | 0.759 | 1.000 | 3,253 |
| **b_Dyadic_Lead_Score_Diff_std** | -0.099 | -0.184 | -0.018 | 0.991 | 1.000 | 3,284 |
| b_Dyadic_Weight_Diff_std | 0.001 | -0.088 | 0.086 | 0.505 | 1.000 | 2,680 |
| b_Nearest_NeighborsnonMnearest | 0.247 | -0.017 | 0.515 | 0.965 | 1.001 | 2,438 |
| b_Chorus_Shapeexploded | 0.189 | -0.278 | 0.682 | 0.784 | 1.001 | 3,022 |
| b_Chorus_Shapenormal | 0.245 | -0.147 | 0.624 | 0.887 | 1.000 | 2,570 |
| b_Effective_Chorus_Size_std | 0.034 | -0.036 | 0.110 | 0.821 | 1.000 | 3,476 |
| **b_Instantaneous_Chorus_Size_std** | 0.213 | 0.124 | 0.298 | 1.000 | 1.001 | 3,213 |
| **b_Ref_Male_Int_Call_Period_std** | 0.177 | 0.020 | 0.335 | 0.986 | 1.001 | 2,742 |
| b_Asynchrony_Run_Dur_std | -0.007 | -0.163 | 0.152 | 0.534 | 1.000 | 2,696 |
| b_Run_Start_Time_Known1 | 0.059 | -0.118 | 0.212 | 0.759 | 1.000 | 6,094 |
| **b_Dyadic_Int_Call_Period_Diff_std:Previous_Onset_Delay_std** | -0.621 | -0.729 | -0.521 | 1.000 | 1.002 | 1,650 |
| b_Nearest_NeighborsnonMnearest:Chorus_Shapeexploded | -0.054 | -0.484 | 0.388 | 0.596 | 1.001 | 3,056 |
| b_Nearest_NeighborsnonMnearest:Chorus_Shapenormal | -0.168 | -0.489 | 0.165 | 0.841 | 1.000 | 2,384 |
| b_Effective_Chorus_Size_std:Instantaneous_Chorus_Size_std | 0.015 | -0.041 | 0.079 | 0.686 | 1.000 | 3,218 |
| b_Asynchrony_Run_Dur_std:Run_Start_Time_Known1 | 0.140 | -0.025 | 0.313 | 0.948 | 1.002 | 3,191 |
| bs_sDyadic_Int_Call_Period_Diff_std_1 | 0.142 | -0.817 | 1.112 | 0.618 | 1.001 | 5,328 |
| bs_sPrevious_Onset_Delay_std_1 | -0.213 | -1.213 | 0.781 | 0.665 | 1.000 | 8,355 |
| ***Group-Level (Random) Effects (SD)*** | | | | | | |
| sd_Chorus_Name__Intercept | 0.172 | 0.001 | 0.354 | 1.000 | 1.001 | 642 |
| sd_Chorus_Name:Dyad_ID__Intercept | 0.267 | 0.096 | 0.416 | 1.000 | 1.014 | 303 |
| sd_Chorus_Name:Dyad_ID__Previous_Onset_Delay_std | 0.620 | 0.522 | 0.724 | 1.000 | 1.000 | 2,106 |
| sd_Chorus_Name:Dyad_ID__Effective_Chorus_Size_std | 0.221 | 0.129 | 0.304 | 1.000 | 1.001 | 1,375 |
| sd_Chorus_Name:Dyad_ID__Instantaneous_Chorus_Size_std | 0.157 | 0.001 | 0.271 | 1.000 | 1.006 | 437 |
| sd_Chorus_Name:Dyad_ID__Asynchrony_Run_Dur_std | 0.109 | 0.000 | 0.246 | 1.000 | 1.004 | 714 |
| sd_mmRef_Male_IDOtr_Male_ID__Intercept | 0.400 | 0.091 | 0.622 | 1.000 | 1.010 | 323 |
| ***Smooth Term Variances (Wiggliness)*** | | | | | | |
| sds_sDyadic_Int_Call_Period_Diff_std_1 | 2.783 | 1.025 | 5.275 | 1.000 | 1.002 | 1,464 |
| sds_sPrevious_Onset_Delay_std_1 | 17.677 | 9.972 | 30.540 | 1.000 | 1.003 | 804 |

**Table S4**. Medians and upper and lower 95% *HDI*s of posterior distributions of fixed effects, random effects, and smooth terms from the Synchrony_Initiation_Model. Also shown are Probability of Direction (*pd*), R-hat, and effective sample size. Bolded fixed effects were robust predictors (*pd* > 97.5%).

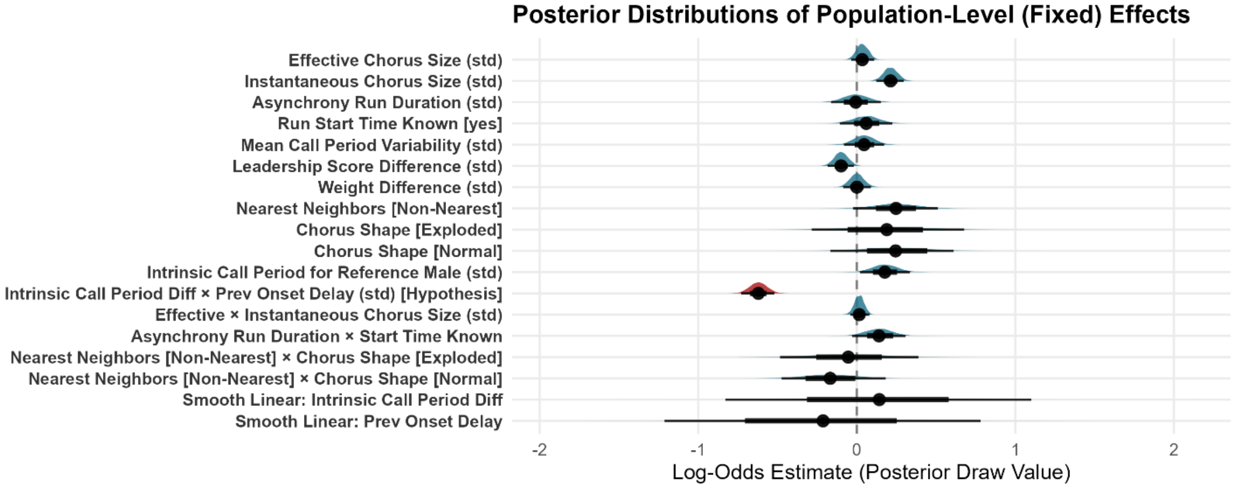


**FIG. S3**. Posterior distributions of population-level (fixed) effects on dyadic synchrony initiation probability (***Synchrony_Initiation_Model***). Distributions are posterior parameter estimates (on the log-odds scale) derived from the Bayesian generalized additive mixed model (GAMM). Density curves depict the full posterior probability density, points denote the posterior medians, and horizontal bars indicate the nested 66% and 95% Highest Density Intervals (*HDI*s). The linear interaction representing the Temporal Offsetting Hypothesis is highlighted in red; as can be seen it is a highly robust and certain predictor of synchrony initiation probability. The parameters distributions presented here with regard to the non-linear smooths are only the linear portions and so can not be directly interpreted. However, the robustness of smooth parameters can be evaluated in the coefficient table above and the variance component plot below, and the conditional marginal effects of smooth terms are visualized in Fig. 2C, in-text.

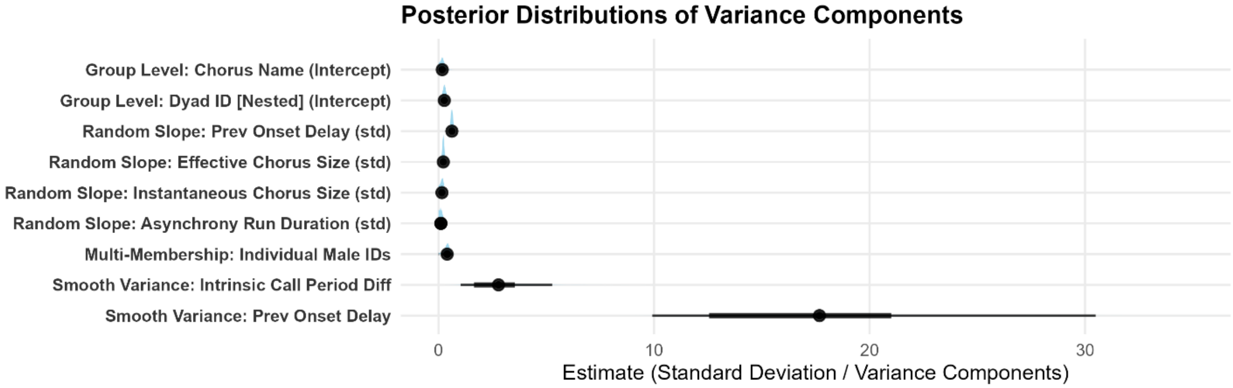


**FIG. S4**. Posterior distributions of group-level (random) effects and smooth term variances for dyadic synchrony initiation probability (Synchrony_Initiation_Model). Distributions represent posterior parameter estimates (on the standard deviation scale) derived from the Bayesian generalized additive mixed model (GAMM). Density curves depict the full posterior probability density, points denote the posterior medians, and horizontal bars indicate the nested 66% and 95% Highest Density Intervals (*HDI*s). These variance components quantify the structural variation in baseline probability and slopes across sampling units (choruses and nested dyads) as well as the magnitude of "wiggliness" (spline complexity) allowed for the non-linear smooth terms.

***SYNCHRONY PERSISTENCE MODEL***

This model investigated the factors influencing the probability that dyads calling synchronously persisted in calling synchronously into the next call cycle. Thus, each row of the dataframe represented a dyad *whose most recent calls had been produced synchronously*, and the outcome variable was a binary response denoting whether their next calls occurred in synchrony (1; synchrony persisted) or not (0). For this model, n = 2,744 call cycles involving 207 unique dyads (3 dyads never synchronized and so are excluded here), in 14 choruses. We describe the variables included in this model, present the model syntax, and present model results, below.

- *Code for this model, model checks, and model results in RMarkdown file 'GAMM_MODELS_FINALIZED.rmd.*
- *Code for generating visualizations of model results in RMarkdown file 'GAMM_VIZ_FINALIZED.rmd.*

***Modeling Approach:*** **Synchrony Persistence Model**

Here, we included most of the same predictors as in the *Synchrony_Inititation_Model*: *Intrinsic Call Period Difference* (non-linear smooth with 10 knots), *Leadership Score Difference*, *Weight Difference*, *Nearest Neighbors* and *Chorus Shape* and their interaction*, Instantaneous Chorus Size* and *Effective Chorus Size* and their interaction, the current duration of the current synchrony run (*Synchrony Run Duration*; analogous to *Asynchrony Run Duration*, but now the duration of the current synchrony run), and the *Intrinsic Call Period of the Reference Male*. We initially included *Run Start Time Known* and its interaction with synchrony run duration, but we removed this interaction and removed *Run Start Time* as there was too little data for unknown run start times. Additionally, we did not include *Previous Onset Delay* as a predictor in this model as dyads were only included in this model when already calling synchronously, thereby they by definition had absolute *Prior Onset Delays* of <0.1s. We also included *Chorus Stability* as a predictor here (defined below), as we predicted that synchrony persistence would be more probable when the same broad chorusing dynamics that led to synchrony in the previous call cycle persisted into the next call cycle (model syntax in Protocol S3, results in Table S5).

- *Chorus Stability*; This is the cosine similarity between: *i)* the vector of delays from the onset time of the reference male's previous call to the onset times of the calls by other chorus members that occurred in the previous call cycle, and *ii)* the vector of these delays in the current call cycle leading up to the interaction of interest. This can best be thought of as a measure of how similar the sequence of other chorus-mates' calls was in the previous cycle and current cycle (1 = identical); *i.e.*, how stable were the chorusing dynamics from one cycle to the next. This variable was only included in the synchrony persistence model in which dyad members both experienced identical inter-call dynamics due to their call cycles being perfectly aligned while calling synchronously.

```
#Regularizing, weakly informative, priors
custom_priors_final <- c(
    # Fixed Effect Prior
        set_prior("normal(0, 0.5)", class = "b"),
    # Intercept Prior
        set_prior("normal(0, 3)", class = "Intercept"),
    # Random Effect SDs Prior
        set_prior("exponential(1)", class = "sd"),
    # Smooth Term SDs Prior
        set_prior("cauchy(0, 0.5)", class = "sds"))

#Model
Synchrony_Persistence_Model <- brm(synchrony_persistence_event ~
    #Test of Temporal Offsetting Hypothesis
            s(Intrinsic_Call_Period_Difference_std, k = 10) +
    #Dyadic Control Variables
            Mean_Call_Period_Variability_std +
            Leadership_Score_Difference_std +
            Weight_Difference_std +
            Nearest_Neighbors * Chorus_Shape +
    #Broader Chorus Dynamics Control Variables
           Effective_Chorus_Size_std * Instantaneous_Chorus_Size_std +
           Chorus_Stability_std +
    #Temporal Control Variables
            Synchrony_Run_Duration_std +
            Intrinsic_Call_Period_of_Reference_Male_std +
    #random effects
           (1 | Chorus_Name/Dyad_ID) +                              #dyad_ID nested in chorus_ID
           (0 + Effective_Chorus_Size_std +
             Instantaneous_Chorus_Size_std +
             Chorus_Stability_std || Chorus_Name : Dyad_ID) + #uncorrelated random slopes
           (1 | mm(Reference_Male_ID, Other_Male_ID)),        #mulitmembership term for individual contributions
    #Run settings
  data = data,
  family = bernoulli(link = "cloglog"),
  prior = custom_priors_final,
  init=0.1,                                                 #restrict initial values to stabilize cloglog link initialization
  chains = 4,
  cores = 4,
  threads = threading(2),
  iter = 4500,
  warmup = 2000,
  refresh = 5,
  backend = "cmdstanr",
  control = list(
    adapt_delta = 0.999,                                    # small step size to eliminate divergent transitions
    max_treedepth = 15                                      # Deeper tree exploration
))
```

**Protocol S2**. Brms syntax for Synchrony_Persistence_Model.

#### *Model Results: Synchrony_Persistence_Model*

| Parameter | Median | 95% HDI Lower | 95% HDI Upper | pd | R-hat | ESS |
|---|---|---|---|---|---|---|
| ***Population-Level (Fixed) Effects*** | | | | | | |
| **b_Intercept** | -1.370 | -1.746 | -1.010 | 1.000 | 1.000 | 7,707 |
| b_Dyadic_Call_Period_Var_std | -0.021 | -0.189 | 0.150 | 0.597 | 1.000 | 8,415 |
| b_Dyadic_Lead_Score_Diff_std | 0.067 | -0.048 | 0.179 | 0.873 | 1.001 | 8,482 |
| b_Dyadic_Weight_Diff_std | 0.062 | -0.055 | 0.177 | 0.849 | 1.000 | 8,785 |
| b_Nearest_NeighborsnonMnearest | 0.213 | -0.144 | 0.565 | 0.883 | 1.000 | 7,608 |
| b_Chorus_Shapeexploded | 0.145 | -0.408 | 0.654 | 0.703 | 1.001 | 8,444 |
| b_Chorus_Shapenormal | 0.032 | -0.388 | 0.456 | 0.562 | 1.000 | 7,819 |
| **b_Effective_Chorus_Size_std** | 0.117 | 0.007 | 0.226 | 0.982 | 1.000 | 10,396 |
| b_Instantaneous_Chorus_Size_std | -0.120 | -0.243 | 0.004 | 0.970 | 1.000 | 9,530 |
| b_Ref_Male_Int_Call_Period_std | 0.111 | -0.093 | 0.304 | 0.868 | 1.001 | 8,295 |
| b_Synchrony_Run_Dur_std | -0.016 | -0.087 | 0.054 | 0.669 | 1.000 | 9,701 |
| b_Chorus_Stability_std | 0.043 | -0.041 | 0.130 | 0.848 | 1.001 | 11,179 |
| b_Nearest_NeighborsnonMnearest:Chorus_Shapeexploded | 0.184 | -0.335 | 0.736 | 0.746 | 1.001 | 8,523 |
| b_Nearest_NeighborsnonMnearest:Chorus_Shapenormal | 0.106 | -0.319 | 0.548 | 0.687 | 1.000 | 7,345 |
| **b_Effective_Chorus_Size_std:Instantaneous_Chorus_Size_std** | -0.179 | -0.263 | -0.086 | 1.000 | 1.001 | 10,718 |
| bs_sDyadic_Int_Call_Period_Diff_std_1 | -0.002 | -1.000 | 0.918 | 0.502 | 1.000 | 18,608 |
| ***Group-Level (Random) Effects (SD)*** | | | | | | |
| sd_Chorus_Name__Intercept | 0.084 | 0.000 | 0.250 | 1.000 | 1.001 | 4,051 |
| sd_Chorus_Name:Dyad_ID__Intercept | 0.471 | 0.340 | 0.621 | 1.000 | 1.000 | 3,109 |
| sd_Chorus_Name:Dyad_ID__Effective_Chorus_Size_std | 0.392 | 0.257 | 0.539 | 1.000 | 1.001 | 2,688 |
| sd_Chorus_Name:Dyad_ID__Instantaneous_Chorus_Size_std | 0.349 | 0.179 | 0.516 | 1.000 | 1.001 | 1,617 |
| sd_Chorus_Name:Dyad_ID__Chorus_Stability_std | 0.223 | 0.042 | 0.369 | 1.000 | 1.002 | 1,762 |
| sd_mmRef_Male_IDOtr_Male_ID__Intercept | 0.168 | 0.000 | 0.430 | 1.000 | 1.002 | 1,583 |
| ***Smooth Term Variances (Wiggliness)*** | | | | | | |
| sds_sDyadic_Int_Call_Period_Diff_std_1 | 1.877 | 0.867 | 3.720 | 1.000 | 1.001 | 6,447 |

**Table S5.** Medians and upper and lower 95% *HDI*s of posterior distributions of fixed effects, random effects, and smooth terms from the Synchrony_Persistence_Model. Also shown are Probability of Direction (*pd*), R-hat, and effective sample size. Bolded fixed effects were robust predictors (*pd* > 97.5%).

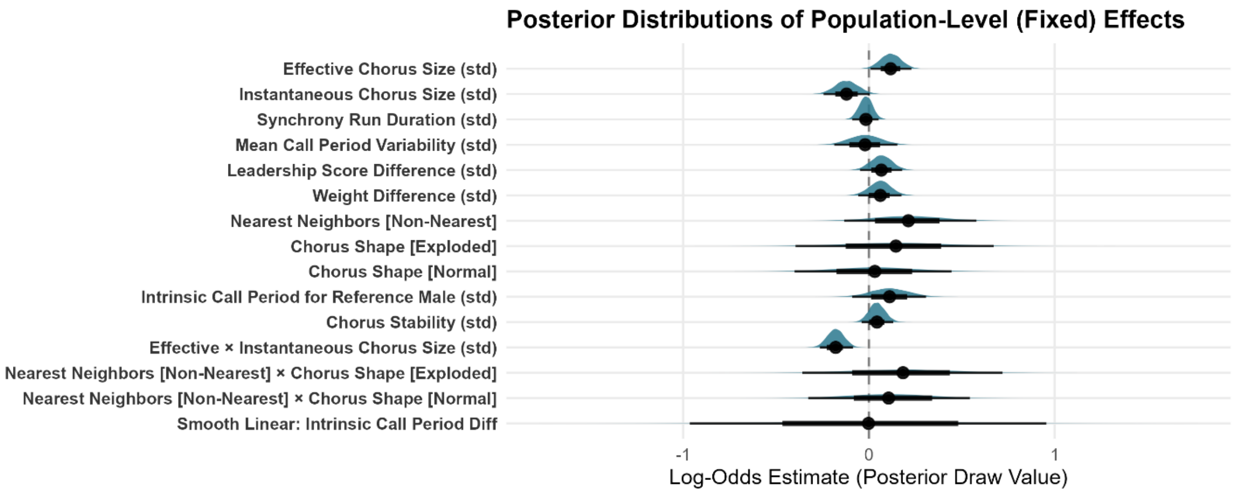


**FIG. S5**. Posterior distributions of population-level (fixed) effects on dyadic synchrony persistence probability (***Synchrony_Persistence_Model***). Distributions are posterior parameter estimates (on the log-odds scale) derived from the Bayesian generalized additive mixed model (GAMM). Density curves depict the full posterior probability density, points denote the posterior medians, and horizontal bars indicate the nested 66% and 95% Highest Density Intervals (*HDI*s). The parameters distributions presented here with regard to the non-linear smooths are only the linear portions and so can not be directly interpreted. However, the robustness of smooth parameters can be evaluated in the coefficient table above and the variance component plot below, and the conditional marginal effects of smooth terms are visualized in Fig. 2C, in-text.

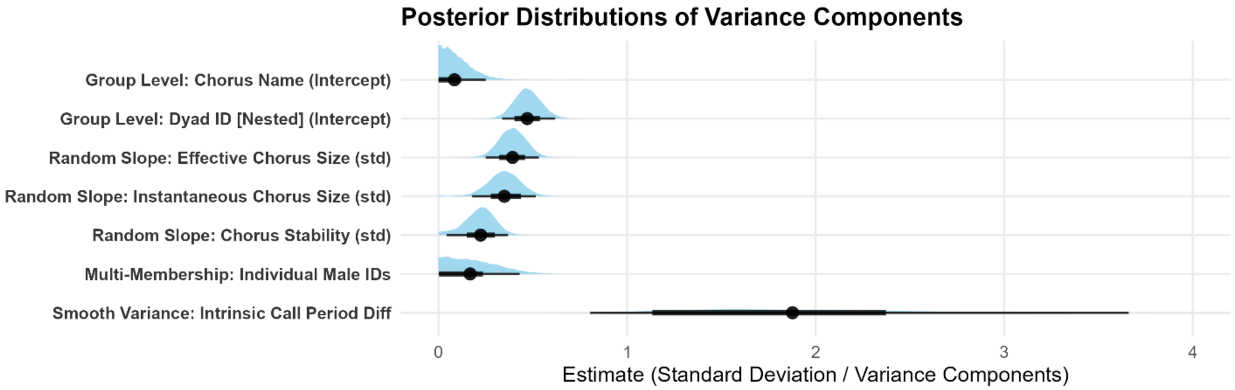


**FIG. S6**. Posterior distributions of group-level (random) effects and smooth term variances for dyadic synchrony initiation probability (Synchrony_Persistence_Model). Distributions represent posterior parameter estimates (on the standard deviation scale) derived from the Bayesian generalized additive mixed model (GAMM). Density curves depict the full posterior probability density, points denote the posterior medians, and horizontal bars indicate the nested 66% and 95% Highest Density Intervals (*HDI*s). These variance components quantify the structural variation in baseline probability and slopes across sampling units (choruses and nested dyads) as well as the magnitude of "wiggliness" (spline complexity) allowed for the non-linear smooth terms.

**Supplementary Text S7. The Influence of Dyadic Call Period Differences on Onset Delay Variability and Occupation of 'Offset States'**

In addition to larger call-period differences influencing per-call synchrony onset probabilities due to the Temporal Offsetting phenomenon described in-text, larger call period differences will generate phase mismatches among dyad members that may lead to more variable dyadic call-timing associations (onset delays). This could influence the proportion of time dyads spend in offset vs. additive states, and how often they transition between these states. Thus, we investigated these relationships. For these analyses, we used the absolute value of dyads' intrinsic call period differences as predictor variables. This is because we would expect that the same magnitude of call period deviation in any direction to generate identical phase mismatches that would increase the variability of dyadic onset delays to a similar degree. We then calculated three metrics from dyads when they were engaged in runs of asynchronous calling (n = 210 dyads for each model): ***1)*** the autocorrelation coefficient of their onset delays at a lag of 1 call cycle (i.e. how strongly correlated were the onset delays in one call cycle with the onset delays in the cycle immediately following it; Protocol S4); ***2)*** the total count of times that a dyad transitioned between being in an offset state to an additive state (Protocol S5), and ***3)*** the total count of call cycles spent in offset states vs. additive states (Protocol S6). We also controlled for dyad members' Mean_Call_Period_Variability scores (described in Supplementary Text S6), as the intrinsically variability of dyad members would also logically influence these outcomes.

While controlling for dyad-members' intrinsic call period variabilities, greater absolute call period differences robustly decreased the cycle-to-cycle autocorrelation of dyadic onset delays during asynchronous calling (i.e., increased cycle-to-cycle onset delay variability) (B-LMM: $\beta_{median}$ = -0.11, 95% *HDI* = [-0.13, -0.08], *pd* = **100%**), and increased the rate at which dyads transitioned between offset and additive states (B-GAMM: $\beta_{median}$ = 0.27, 95% *HDI* = [0.22, 0.31], *pd* = **100%**). However, call period difference did not robustly influence the proportion of call cycles dyads spent in an offset state (B-GLMM: $\beta_{median}$ = -0.02, 95% *HDI* = [-0.08, 0.03], *pd* = 80%). This suggests that the positive effect of call period difference on overall synchrony rates seen in our study primarily arises because dyads with larger call-period differences experience higher synchrony initiation probabilities while in offset states due to the temporal offsetting phenomenon (see in-text results), rather than occupying these states more often. However, there may also be more subtle effects of the more variable call-timing relationships in dyads with larger call period differences that might contribute to their higher synchrony run initiation rates.

- *Code for these models, model checks, and model results in RMarkdown file 'DYADIC_SYNCH_OUTCOMES_FINALIZED.rmd'.*

***MODEL 1***

```
lag1_AutoCorrelation_Coef ~ standardized(abs(Intrinsic_Call_Period_Difference)) +
                              standardized(Mean_Call_Period_Variability) +
                              (1 | Chorus_Name) +
                              (1 | multimembership(Reference_Male_ID, Other_Male_ID)
```

**Protocol S4**. Brms syntax for model with the autocorrelation coefficient of their onset delays at a lag of 1 call cycle as the response variable (family = Gaussian).

***Results***

| Parameter | Median | 95% HDI Lower | 95% HDI Upper | pd | R-hat | ESS |
|---|---|---|---|---|---|---|
| ***Population-Level (Fixed) Effects*** | | | | | | |
| b_Intercept | 0.271 | 0.247 | 0.296 | 1.000 | 1.001 | 3,047 |
| b_Abs_ICP_Difference_std | -0.106 | -0.130 | -0.080 | 1.000 | 1.000 | 3,613 |
| b_Dyadic_Call_Period_Var_std | -0.069 | -0.094 | -0.044 | 1.000 | 1.001 | 3,469 |
| ***Group-Level (Random) Effects (SD)*** | | | | | | |
| sd_Chorus_Name__Intercept | 0.016 | 0.000 | 0.044 | 1.000 | 1.002 | 1,345 |
| sd_mmRef_Male_IDOtr_Male_ID__Intercept | 0.053 | 0.000 | 0.099 | 1.000 | 1.012 | 501 |
| ***Distributional Parameters (Residual Noise)*** | | | | | | |
| sigma | 0.132 | 0.116 | 0.147 | 1.000 | 1.003 | 1,417 |

**Table S6.** Medians and upper and lower 95% *HDI*s of posterior distributions of fixed effects, random effects, and smooth terms. Also shown are Probability of Direction (*pd*), R-hat, and effective sample size.

***MODEL 2***

Count_Transitions_Between_States ~ standardized(abs(Intrinsic_Call_Period_Difference)) +
s(standardized(Mean_Call_Period_Variability), k = 5) +
offset(log(Total_Possible_Transitions)) +
(1 | Chorus_Name) +
(1 | multimembership(Reference_Male_ID, Other_Male_ID)

**Protocol S5**. Brms syntax for model with the total count of transitions between offset and additive states as the response variable (family = Poisson; GAMM as diagnostics indicated Mean_Call_Period_Variability needed a basis function).

***Results***

| Parameter | Median | 95% HDI Lower | 95% HDI Upper | pd | R-hat | ESS |
|---|---|---|---|---|---|---|
| ***Population-Level (Fixed) Effects*** | | | | | | |
| b_Intercept | -1.677 | -1.748 | -1.602 | 1.000 | 1.002 | 2,567 |
| b_Abs_ICP_Difference_std | 0.265 | 0.225 | 0.311 | 1.000 | 1.001 | 4,367 |
| bs_sDyadic_Call_Period_Var_std_1 | 4.121 | 2.124 | 6.318 | 1.000 | 1.000 | 3,398 |
| ***Group-Level (Random) Effects (SD)*** | | | | | | |
| sd_Chorus_Name__Intercept | 0.099 | 0.011 | 0.183 | 1.000 | 1.006 | 835 |
| sd_mmRef_Male_IDOtr_Male_ID__Intercept | 0.139 | 0.019 | 0.228 | 1.000 | 1.003 | 699 |
| ***Smooth Term Variances (Wiggliness)*** | | | | | | |
| sds_sDyadic_Call_Period_Var_std_1 | 2.727 | 1.029 | 5.859 | 1.000 | 1.002 | 1,816 |

**Table S7**. Medians and upper and lower 95% *HDI*s of posterior distributions of fixed effects, random effects, and smooth terms. Also shown are Probability of Direction (*pd*), R-hat, and effective sample size.

***MODEL 3***

```
Count_Calls_in_Offset_State ~ standardized(abs(Intrinsic_Call_Period_Difference)) +
                              standardized(Mean_Call_Period_Variability) +
                              offset(log(Total_Possible_Calls)) +
                              (1 | Chorus_Name) +
                              (1 | multimembership(Reference_Male_ID, Other_Male_ID)
```

**Protocol S6**. Brms syntax for model with the total count of call cycles in an offset state as the response variable (family = negative binomial).

***Results***

| Parameter | Median | 95% HDI Lower | 95% HDI Upper | pd | R-hat | ESS |
|---|---|---|---|---|---|---|
| ***Population-Level (Fixed) Effects*** | | | | | | |
| b_Intercept | -0.671 | -0.716 | -0.628 | 1.000 | 1.001 | 4,928 |
| b_Dyadic_Call_Period_Var_std | -0.022 | -0.073 | 0.028 | 0.804 | 1.000 | 4,083 |
| b_Abs_ICP_Difference_std | 0.001 | -0.049 | 0.051 | 0.518 | 1.001 | 4,316 |
| ***Group-Level (Random) Effects (SD)*** | | | | | | |
| sd_Chorus_Name__Intercept | 0.019 | 0.000 | 0.058 | 1.000 | 1.001 | 1,969 |
| sd_mmRef_Male_IDOtr_Male_ID__Intercept | 0.039 | 0.000 | 0.108 | 1.000 | 1.002 | 1,750 |
| ***Distributional Parameters (Residual Noise / Dispersion)*** | | | | | | |
| shape | 15.615 | 12.097 | 19.871 | 1.000 | 1.001 | 6,585 |

**Table S8**. Medians and upper and lower 95% *HDI*s of posterior distributions of fixed effects, random effects, and smooth terms. Also shown are Probability of Direction (*pd*), R-hat, and effective sample size.

**Supplementary Text S8. Clearly Demonstrating that Temporal Offsetting Drives the Positive Relationship Between Dyadic Call Period Differences and Synchrony Initiation Probability**

We tested our Temporal Offsetting Hypothesis by including a linear interaction in our *Synchrony_Initiation_Model* GAMM between intrinsic call period difference and previous onset delay. This interaction was a robust predictor of synchrony onset probability (B-GAMM: $\beta_{median}$ = -0.62, 95% *HDI*: [-0.73, -0.52], *pd* = **100%**; Table S4; Fig. S3), and removal of this interaction diminished marginal $R^2$ by 40% and substantially decreased model predictive power ($\Delta elpd$ = 14.9, $se_{diff}$ = 6, $\Delta elpd/se_{diff}$ = **2.48**). These results strongly support Temporal Offsetting as an important mechanism driving synchrony initiation.

Furthermore, a corollary of this hypothesis was that the Temporal Offsetting phenomenon was a key mechanism leading to higher synchrony initiation probabilities for dyads with more dissimilar call periods. This was because these larger call period differences might be more effectively offset by the onset delays that correspond to the typical asynchronous calling interactions this species engages in (~0.3s and ~0.6s for overlap and alternation, respectively; Fig1A, in-text). Thus, the linear interaction between intrinsic call period difference and previous onset delay should be a key mediator of this relationship between larger call period differences and synchrony initiation probability in our model. However, demonstrating that this interaction itself was the key mechanism driving this relationship was challenging. For instance, visualizations of the conditional marginal effect of call period difference on synchrony initiation probability looked nearly identical in the full model (including this key interaction), and in the reduced model (excluded the interaction). While initially counterintuitive, this pattern arises because when the interaction term is removed, the constituent main effects remain in the model and can absorb the explanatory power of that interaction. In other words, the reduced model still 'knows' that dyads with more dissimilar intrinsic call periods have higher synchrony initiation probabilities, even though it no longer 'knows' the underlying structural reason why. Consequently, the model's calculation of the overall marginal effect of dyadic call period difference when all other variables are held constant remains largely unchanged.

To isolate and visually demonstrate that Temporal Offsetting itself drives this relationship, we calculated the Area Under the Curve (AUC) across our full range of prior onset delays, evaluated at 50 distinct points across the range of dyadic call period differences. Because dyadic call period difference was calculated as the intrinsic call period of the reference male minus that of the other male in the dyad, it could be positive or negative depending on which male had the longer intrinsic call period. Similarly, previous onset delays could be positive or negative depending on which male called first in the preceding call cycle. Thus, if temporal offsetting is the mechanism leading dyads with more dissimilar call periods to initiate synchrony at higher rates, dyads with larger call period differences should *only* have higher synchrony initiation probabilities when in ‘offset states’ in which previous onset delay and dyadic call period differences carry opposite signs (+/- or -/+, not +/+ or -/-). To visually reveal this, we plotted the AUC values corresponding to positive and negative previous onset delays separately across our entire range of intrinsic call period differences (Fig. 2D, in-text). As expected, larger dyadic call period differences only increased synchrony initiation probabilities when dyads were in offset states. This is shown in the figure by the fact that larger negative dyadic call period differences only had elevated synchrony probabilities when experiencing positive onset delays, and vice versa; i.e. when in offset states. This provides strong evidence that temporal offsetting drives the observed positive relationship between dyadic call period difference and synchrony initiation probabilities, which are a key driver of overall synchrony rate (see Supplementary Text S9).

- *Code for this analytical pipeline and results visualization in RMarkdown file ‘GAMM_VIZ_FINALIZED.rmd’.*

**Supplementary Text S9. Drivers of Overall Dyadic Synchrony Rate**

To investigate whether dyads’ rate of synchrony run initiations or their typical durations better predicted total dyadic synchronous call counts, we constructed linear mixed-effects models (LMMs) in brms. We modeled $\log_{10}$(total synchronous calls) as a function of $\log_{10}$(total synchrony runs) in one model (Protocol S7), and as a function of $\log_{10}$(mean synchrony run duration) in another (Protocol S8). Inclusion of an offset here was not necessary as both sides of the equation will be similarly limited by the overall call rates of the dyad members. For these models, both the response and predictors were log-transformed to reflect the underlying multiplicative relationship between synchrony run initiation rate, run duration, and total count of synchronous calls (Total Synchronous Calls = Synchrony Run Rate * Synchrony Run Duration), thereby linearizing the model and making these relationships additive. These log-transformations allowed these relationships to be captured as scale-free proportional changes, with standardization [(x-mean(x))/SD(x)] then allowing direct comparisons of effect sizes across models. Dyads that produced zero synchronous calls did not exhibit observable synchrony run durations. Thus, we opted to only include dyads here that produced at least 1 synchronous call (n = 207 of 210 total dyads). Model results are presented below as well as a model comparison demonstrating that the model including synchrony run initiation rate had substantially superior predictive power when explaining dyads' total counts of synchronous calls (Table S11). This suggests that variation in synchrony run initiation rate was the main driver of variation in dyadic synchrony rates.

- *Code for these models, model checks, and model results in RMarkdown file 'DYADIC_SYNCH_OUTCOMES_FINALIZED.rmd'.*

**RATE_MODEL**

**Formula:** log10(Total_Synchronous_Calls) ~ standardized(log10(Synchrony_Run_Initiation_Rate)) + (1 | Chorus_Name)

**Protocol S7**. Brms syntax for model with Synchrony_Run_Initiation_Rate as the predictor variable (family = Gaussian).

***Results:***

| Parameter | Median | 95% HDI Lower | 95% HDI Upper | pd | R-hat | ESS |
|---|---|---|---|---|---|---|
| *Population-Level (Fixed) Effects* | | | | | | |
| b_Intercept | 2.423 | 2.351 | 2.499 | 1.000 | 1.003 | 1,630 |
| b_log_freq_std | 0.512 | 0.466 | 0.561 | 1.000 | 1.001 | 4,465 |
| *Group-Level (Random) Effects (SD)* | | | | | | |
| sd_Chorus_Name__Intercept | 0.097 | 0.016 | 0.181 | 1.000 | 1.001 | 986 |
| *Distributional Parameters (Residual Noise / Dispersion)* | | | | | | |
| sigma | 0.340 | 0.308 | 0.376 | 1.000 | 1.001 | 2,866 |

**Table S9**. Medians and upper and lower 95% *HDI*s of posterior distributions of fixed effects, random effects, and smooth terms. Also shown are Probability of Direction (*pd*), R-hat, and effective sample size.

**DURATION_MODEL**

log10(Total_Synchronous_Calls) ~ standardize(log10(Mean_Synchrony_Run_Duration)) + (1 | Chorus_Name)

**Protocol S8**. Brms syntax for model with Mean_Synchrony_Run_Duration as the predictor variable (family = Gaussian)..

***Results:***

| Parameter | Median | 95% HDI Lower | 95% HDI Upper | pd | R-hat | ESS |
|---|---|---|---|---|---|---|
| ***Population-Level (Fixed) Effects*** | | | | | | |
| b_Intercept | 2.422 | 2.324 | 2.519 | 1.000 | 1.001 | 3,530 |
| b_log_dur_std | 0.225 | 0.151 | 0.305 | 1.000 | 1.000 | 4,852 |
| ***Group-Level (Random) Effects (SD)*** | | | | | | |
| sd_Chorus_Name__Intercept | 0.089 | 0.000 | 0.208 | 1.000 | 1.001 | 1,372 |
| ***Distributional Parameters (Residual Noise / Dispersion)*** | | | | | | |
| sigma | 0.573 | 0.514 | 0.631 | 1.000 | 1.000 | 4,564 |

**Table S10**. Medians and upper and lower 95% *HDI*s of posterior distributions of fixed effects, random effects, and smooth terms. Also shown are Probability of Direction (*pd*), R-hat, and effective sample size.

***Model Comparison Table***

| Predictor Model Structure | Δ elpd[1] | se (diff) | elpd_loo | se (elpd) | LOOIC |
|---|---|---|---|---|---|
| Synchrony Run Frequency Model | 0.00 | 0.00 | -76.27 | 13.13 | 152.55 |
| Synchrony Run Duration Model | -105.64 | 16.66 | -181.91 | 12.13 | 363.83 |

[1]The top-performing model is automatically anchored at Δ elpd = 0.00. Deficits where Δ elpd / se < 2.00 represent highly competitive alternative structures.

**Table S11**. LOO-IC model comparison table for comparing the predictive importance of dyadic synchrony run initiation rate and duration in predicting overall dyadic synchrony rates.

**Supplementary Text S10. Intrinsic Call Period as a Driver of Variation in Individual Synchrony Rates**

We had previously found that larger call period differences among dyad members increased dyadic synchrony rates (Fig. 2A, in-text). Thus, we predicted that males with extreme call periods would engage in synchrony at overall higher rates as they would tend to exhibit larger call-period differences with the other males in their choruses. However, it may not simply be how long a male's call period is in absolute terms that is most important, but rather how different their call period is from the other males in their chorus. Thus, we also calculated more chorus-specific call period deviation scores such as: 1) the difference between a male's intrinsic call period and the median intrinsic call period of their chorus (difference_from_chorus_median); 2) the mean absolute difference between a male's intrinsic call period and those of his chorus-mates (mean_absolute_call_period_difference); 3) the count of chorus-mates that a male's intrinsic call period differed from by more than 0.1s (count_call_period_different). We then constructed negative binomial models that regressed males' aggregate synchrony counts on each of these metrics, including an offset for his total number of calls. Results are presented below.

- *Code for these models, model checks, and model results in RMarkdown file 'INDIVIDUAL_SYNCH_OUTCOMES_FINALIZED.rmd'.*

***Regression on Intrinsic Call Period***

| Parameter | Median | 95% HDI Lower | 95% HDI Upper | pd | R-hat | ESS |
|---|---|---|---|---|---|---|
| ***Population-Level (Fixed) Effects*** | | | | | | |
| b_Intercept | -0.728 | -0.788 | -0.669 | 1.000 | 1.002 | 2,353 |
| b_Intrinsic_Call_Period_std | 0.098 | 0.051 | 0.149 | 1.000 | 1.001 | 4,222 |
| ***Group-Level (Random) Effects (SD)*** | | | | | | |
| sd_Chorus_Name__Intercept | 0.057 | 0.000 | 0.127 | 1.000 | 1.001 | 1,275 |
| ***Distributional Parameters (Residual Noise / Dispersion)*** | | | | | | |
| shape | 36.061 | 18.581 | 60.378 | 1.000 | 1.001 | 3,863 |

**Table S12**. Medians and upper and lower 95% *HDI*s of posterior distributions of fixed effects, random effects, and smooth terms. Also shown are Probability of Direction (*pd*), R-hat, and effective sample size.

***Regression on difference_from_chorus_median***

| Parameter | Median | 95% HDI Lower | 95% HDI Upper | pd | R-hat | ESS |
|---|---|---|---|---|---|---|
| ***Population-Level (Fixed) Effects*** | | | | | | |
| b_Intercept | -0.728 | -0.790 | -0.660 | 1.000 | 1.000 | 2,681 |
| b_diff_from_group_median_std | 0.081 | 0.035 | 0.131 | 1.000 | 1.000 | 4,817 |
| ***Group-Level (Random) Effects (SD)*** | | | | | | |
| sd_Chorus_Name__Intercept | 0.078 | 0.002 | 0.152 | 1.000 | 1.002 | 692 |
| ***Distributional Parameters (Residual Noise / Dispersion)*** | | | | | | |
| shape | 34.594 | 18.063 | 56.935 | 1.000 | 1.001 | 2,895 |

**Table S13.** Medians and upper and lower 95% *HDI*s of posterior distributions of fixed effects, random effects, and smooth terms. Also shown are Probability of Direction (*pd*), R-hat, and effective sample size.

***Regression on mean_absolute_call_period_difference***

| Parameter | Median | 95% HDI Lower | 95% HDI Upper | pd | R-hat | ESS |
|---|---|---|---|---|---|---|
| ***Population-Level (Fixed) Effects*** | | | | | | |
| b_Intercept | -0.731 | -0.806 | -0.658 | 1.000 | 1.002 | 1,683 |
| b_mean_mate_diff_unsigned_std | 0.104 | 0.055 | 0.154 | 1.000 | 1.001 | 3,411 |
| ***Group-Level (Random) Effects (SD)*** | | | | | | |
| sd_Chorus_Name__Intercept | 0.096 | 0.014 | 0.173 | 1.000 | 1.009 | 983 |
| ***Distributional Parameters (Residual Noise / Dispersion)*** | | | | | | |
| shape | 40.595 | 20.657 | 71.848 | 1.000 | 1.001 | 2,754 |

**Table S14**. Medians and upper and lower 95% *HDI*s of posterior distributions of fixed effects, random effects, and smooth terms. Also shown are Probability of Direction (*pd*), R-hat, and effective sample size.

#### *Regression on count_call_period_different*

| Parameter | Median | 95% HDI Lower | 95% HDI Upper | pd | R-hat | ESS |
|---|---|---|---|---|---|---|
| ***Population-Level (Fixed) Effects*** | | | | | | |
| b_Intercept | -0.730 | -0.812 | -0.650 | 1.000 | 1.002 | 1,599 |
| b_count_mates_distinct_std | 0.091 | 0.035 | 0.151 | 0.999 | 1.000 | 1,886 |
| ***Group-Level (Random) Effects (SD)*** | | | | | | |
| sd_Chorus_Name__Intercept | 0.112 | 0.026 | 0.212 | 1.000 | 1.003 | 726 |
| ***Distributional Parameters (Residual Noise / Dispersion)*** | | | | | | |
| shape | 36.757 | 19.657 | 62.365 | 1.000 | 1.000 | 1,884 |

**Table S15**. Medians and upper and lower 95% *HDI*s of posterior distributions of fixed effects, random effects, and smooth terms. Also shown are Probability of Direction (*pd*), R-hat, and effective sample size.

#### *Model Comparison Results*

| Predictor Model Structure | Δ elpd[1] | se (diff) | elpd_loo | se (elpd) | LOOIC |
|---|---|---|---|---|---|
| Model 2: Localized Chorus-Deviation Metric B | 0.00 | 0.00 | -328.52 | 6.26 | 657.03 |
| Model 0: Absolute Intrinsic Call Period (Median IOI) | -1.75 | 3.77 | -330.27 | 6.19 | 660.54 |
| Model 3: Alternative Rhythmic Control Architecture | -2.91 | 1.69 | -331.43 | 6.03 | 662.85 |
| Model 1: Localized Chorus-Deviation Metric A | -3.11 | 3.38 | -331.62 | 5.74 | 663.24 |

[1]The top-performing model is automatically anchored at Δ elpd = 0.00. While absolute intrinsic call period is the top model, note that deficits where Δ elpd / se < 2.00 represent highly competitive alternative structures.

**Table S16**. LOO-IC model comparison table for comparing the predictive importance of different intrinsic call period deviation metric in predicting individuals' overall synchrony rates.

Though the top performing model had mean_absolute_call_period_difference with chorus-mates as a predictor, none of these models differed robustly from one another in predictive power ($\Delta elpd/se_{diff} < 2.00$ for all comparisons). Thus, the raw intrinsic call period explained synchrony rate as well as more chorus-specific metrics. This suggests that male call periods in this population strongly cluster around typical values. Thus, males with relatively longer call periods are at a population-level disadvantage that can not be easily overcome through strategic choice of chorus. This could increase the strength of selection against these longer call periods (Supplementary Text S14).

### Supplementary Text S11. Transitivity of Synchrony Leadership and Leadership Scores

#### *Transitivity of Leadership*

For each chorus we constructed a lead/follow matrix populated with the count of synchronous interactions each male led with each other male in the chorus, and identified an overall winner for each dyad (the male that led the majority of synchronous interactions). We then counted the total number of non-transitive triads observed across all of our choruses, e.g. those in which A led B, and B led C, but C led A, or some other variation of non-transitivity. We then generated a null distribution of non-transitive triad counts that would be expected if there were no transitivity in propensities to lead, i.e. if the outcome of interactions were purely random. Over 10,000 permutations, we randomly allocated wins to each dyad member with a probability of 0.5

per interaction while retaining the observed number of synchronous interactions engaged in by each dyad. Each iteration, we counted the number of non-transitive triads observed in each permuted matrix. We then calculated a *p-value* as the proportion of permuted non-transitive triad counts that were less than or equal to our observed count. Propensities to lead were significantly transitive (p = 0.0042; Fig. 3A, in-text).

- *Code for this simulation in Python Jupyter notebook 'Transitivity_permutation_FINALIZED.ipynb'.*

***Leadership Score Calculations***

To calculate 'Leadership Scores' which we used as a proxy for relative effector delay duration, we used the same lead/follow matrices mentioned above to calculate David's scores within each chorus, using the method by Gammell et al, (2003) (*33*). David's scores are a commonly used metric in the animal dominance literature, and they do not require extremely steep hierarchies to perform well (*34*). David's scores are continuous cardinal scores, rather than ordinal ranks, meaning that differences between scores are meaningful regarding relative propensities to lead. Thus, they serve as an ideal proxy for relative effector delay durations due to the intuitive connection between this attribute and propensities to lead or follow when males have their calls triggered synchronously (logic described in detail in-text). We generated David's scores using the getDS() function from the 'steepness' R package (*35*). When doing so, we used the 'Dij' method, which applies a correction to make score calculations more robust when dyads differ substantially in their numbers of observed interactions (*33*). As effector delay duration distributions likely vary among choruses, David's scores were standardized within choruses [(x-mean(x))/SD(x)] prior to analysis.

- *Code for this leadership score calculations in Python Jupyter notebook 'Get_David_Scores_FINALIZED.ipynb'.*

**Supplementary Text S12. Investigating the Correlation Between Leadership Score and Intrinsic Call Period**

As we had identified Leadership Score (as a proxy for relative effector delay durations) and intrinsic call period as key sensorimotor variables driving synchrony outcomes (leading/following propensity and synchrony rate, respectively), we wanted to see whether they were correlated with one another. If they were correlated, this could suggest that they are downstream manifestations of some broader sensorimotor tuning phenotype, rather than distinct attributes. When doing so, we also had to account for chorus-level variation in these attributes. To evaluate this relationship while accounting for chorus-level variation in both traits, we implemented a Bayesian multivariate mixed-effects model (B-MMM) in 'brms'(24) that combined two intercept only models (one for each attribute). We specified a shared group-level random intercept for chorus identity [(1|p|chorus_name)] in our sub-models, allowing for the estimation of covariance at the chorus level, and enabled a residual correlation structure [set_rescor(TRUE)]. This parameterization allowed us to assess the individual-level correlation between these attributes, independent of shared chorus-level baseline shifts. We chose this multivariate approach over a standard univariate regression because regressions assume a direction of causality, whereas we were interested in evaluating correlation. The model syntax is presented below.

- *Code for this model, model checks, and model results in RMarkdown file 'INDIVIDUAL_SYNCH_OUTCOMES_FINALIZED.rmd'.*

**MULTIVARIATE_MODEL**

```
#univariate formulas with linked random effects
bf_LR   <- bf(Leadership_Score_std ~ 1 + (1|p| Chorus_Name))
bf_ICP <- bf(Intrinsic_Call_Period_std ~ 1 + (1 |p| Chorus_Name))

# the multivariate model
mod_corr_mixed <- brm(
  bf_LR + bf_ICP + set_rescor(TRUE),
  data = data,
  family = gaussian(),
  cores = 4
)
```

**Protocol S9**. Brms syntax for multivariate model.

***Model Results***

| Relationship Structure | Median (ρ) | 95% HDI Lower | 95% HDI Upper | pd | R-hat | ESS |
|---|---|---|---|---|---|---|
| Residual Correlation (ρ) [Leadership Score × Intrinsic Call Period] | -0.17 | -0.36 | 0.05 | 0.93 | 1.000 | 5,496 |

**Table S17**. Medians and upper and lower 95% *HDI*s of posterior distributions of residual correation. Also shown are Probability of Direction (*pd*), R-hat, and effective sample size.

Leadership score and intrinsic call period were not robustly correlated, though exhibited a suggestive negative trend (B-MMM: $r_{median}$ = -0.17, 95% *HDI* = [-0.37, 0.04], *pd* = 94%; males with shorter call periods tended to have high leadership scores (i.e., shorter effector delays)).

**Supplementary Text S13. Bayesian Bootstrapping and Sensitivity Analysis**

***Bayesian Bootstrapping***

In-text, we describe how we calculated 'Attractiveness Coefficients$_{\text{Synchrony}}$' ($AC_{\text{Synchrony}}$) scores for all males based on their observed synchrony outcomes in our chorus recordings. These represent multipliers applied to males' baseline attractiveness values to account for their expected synchrony rates and roles. To do so, we used Equation 1 (reprinted below) to combine observed male synchrony outcomes (rates and leadership propensities) with female preference values reported in previous experiments (*4*, *15*). However, we did not use observed values directly. Rather, to rigorously account for uncertainty in $AC_{\text{Synchrony}}$ score estimates arising due to sampling effects in our current study and in previous experiments, we used a Bayesian bootstrapping approach to generate a distribution of 100,000 estimates per male.

During each iteration, the probabilities of producing non-synchronous ($P_{\text{non-synchrony}}$), leading synchronous ($P_{\text{lead|synchrony}}$), and following synchronous ($P_{\text{follow|synchrony}}$) calls for each male were drawn jointly from a multivariate Dirichlet distribution shaped by his observed synchrony outcomes in our chorus recordings. Simultaneously, female preference weights for non-synchronous calls ($W_{\text{non-synchrony}}$) and leading and following synchronous calls ($W_{\text{leader|synchrony}}$, $W_{\text{follow|synchrony}}$) were drawn from two beta distributions shaped by the results and sample sizes of previous experiments(*4*, *15*). These Dirichlet and beta distributions were initialized with weakly informative Laplace priors (pseudo-counts of 1) to ensure conservative estimates and account for finite sample sizes. During each iteration, *i)* values drawn from these distributions were combined using Equation 1 to assign each male an $AC_{Synchrony}$ score, *ii)* pairwise differences among the $AC_{Synchrony}$ scores drawn that iteration for males in the same chorus were calculated, and *iii)* differences between each male's score and the median score of his chorus drawn that iteration were calculated. These latter two calculations gave us simulated distributions of differences among males within the same chorus, and between males and their chorus medians. Form these, we could determine whether males differed robustly from one another, and from their chorus median, if the 95% *HDI* of these difference distributions did not contain 0.

- *Code for these Bayesian bootstrapping simulations and sensitivity analyses in RMarkdown file 'INDIVIDUAL_SYNCH_OUTCOMES.rmd'.*

$$\mathbf{AC_{Synchrony}} = (P_{non\text{-}synchrony} * W_{non\text{-}synchrony}) + [(P_{synchrony} * W_{synchrony}) * [(P_{lead|synchrony} * W_{leader|synchrony}) + (P_{follow|synchrony} * W_{follower|synchrony})]]$$

**Equation 1**. $AC_{Synchrony}$ = Attractiveness Coefficient due to Synchrony; $P_{synchronous}$ and $P_{non\text{-}synchronous}$ = the probability that a male produced a synchronous or non-synchronous call, respectively; $P_{lead|synchrony}$ and $P_{follow|synchrony}$ = the probability that, during a synchronous call, a male was a leader or follower, respectively; $W_{non\text{-}synchronous}$ = the probability weight that a female will choose a non-synchronous call over a synchronous one; $W_{leader}$ and $W_{follower}$ = the probability weight that, if a female chooses a synchronous, she will choose the leader or follower, respectively.

***Choice of Previous Studies to Extract Female Choice Estimates From***

There is some disagreement in the túngara frog literature as to how females respond to synchronous vs. non-synchronous calls. Schwartz and Rand (1991) (*5*) found a weak, non-significant preference for non-synchronous calls in túngara frogs, which contrasts with a much stronger, significant preference reported more recently by Legett *et al.* (2019) (*4*) (Table S1). For baseline female preference parameters used in our Bayesian bootstrapping simulations, we elected to use the preference estimates from Legett *et al.* (*4*) We made this choice for three methodological reasons. First, Legett *et al.* (*4*) utilized a substantially larger sample size (n = 40 in Legett *et al.* (*4*) vs. n = 20 in Schwartz and Rand (*5*)). Second, Legett *et al.* (*4*) used recordings of natural túngara frog calls as stimuli, rather than the synthesized calls used in Schwartz and Rand (*5*), maximizing ecological and acoustic validity and generalizability. Third, Schwartz and Rand (*5*) also reported no preference for leading calls during synchrony in túngara frogs (no precedence effect). This contrasts with the strong precedence effect consistently reported for this species in subsequent studies that featured robust sample sizes and used both natural and synthesized stimuli(*14, 15, 16*) (Table S2). Thus, while Schwartz and Rand (*5*) is a foundational study of the effects of call synchrony, their atypical results compared to more recent studies suggest that experimental artefacts may be present due to historical technological limitations when generating convincing synthetic stimulus calls, or from other aspects of their playback setup.

Due to the above reasons, during our Bayesian bootstrapping simulations we opted to use estimates from *Legett et al.* (*4*) for the strength of female preferences for non-synchronous calls (33/40 females choosing non-synchronous calls over synchronous calls (*4*)), and estimates from Larter and Ryan (2024) (*15*) for the strength of female preferences for leading calls (33/35 females choosing leading calls over following calls during synchrony (*15*)). Crucially, to ensure our conclusions were not unduly influenced by our choice of female preference estimates, we also conducted a comprehensive sensitivity analysis (details below). Our simulation results were largely robust as long as females exhibited even slight preferences for non-synchronous calls and leading calls. Female preferences in these directions are the overwhelming pattern seen in alternating frogs (Tables S1 and S2). Overall, this demonstrates the robustness of our approach to the specific baseline preference estimates chosen.

***Sensitivity Analysis of Bayesian Bootstrapping***

To ensure our results were not unduly influenced by the specific female preference estimates obtained from previous experiments, we performed a sensitivity analysis by repeating this simulation process using all possible strengths of female preferences for non-synchronous calls and leading calls. We did so by iterating through all possible results that could have been obtained in these previous experiments(*4*, *15*). This kept the experimental sample size (and thus uncertainty) consistent across these 'alternate universes' but altered the proportion of females that chose one option or the other. During each iteration, we performed the same Bayesian bootstrapping steps outlined in the section above. Then, to evaluate how much our results would have changed if using these counterfactual female preference values, we computed two metrics for each possible combination of preference strengths; 1) *Statistical Discernibility*: the percentage of male pairs within each chorus that differed robustly from one another in their $AC_{Synchrony}$ scores (0 not contained in 95% *HDI* of their $AC_{Synchrony}$ score difference), and 2) *Statistical Stability*: the Spearman's rank correlation between the means of males' $AC_{Synchrony}$ score distributions generated with counterfactual female preference estimates and those generated with the female preference estimates actually obtained in previous experiments (33/40 females choosing non-synchronous calls over synchronous calls (*4*); 33/35 females choosing leading calls over following calls during synchrony (*15*)).

Results are shown in the figures below (Fig. S7 and S8). As can be seen, within the upper right quadrant in which females showed some degree of preference for both non-synchronous calls and leading calls, both metrics were largely stable within choruses and on average. Female preferences for non-synchronous calls and leading calls is the overwhelming pattern seen in alternating frogs (**Tables S1 and S2**). Thus, that male ranks and differentiation are both largely stable in the upper right quadrant suggests our results are qualitatively robust to the precise strength of female preferences.

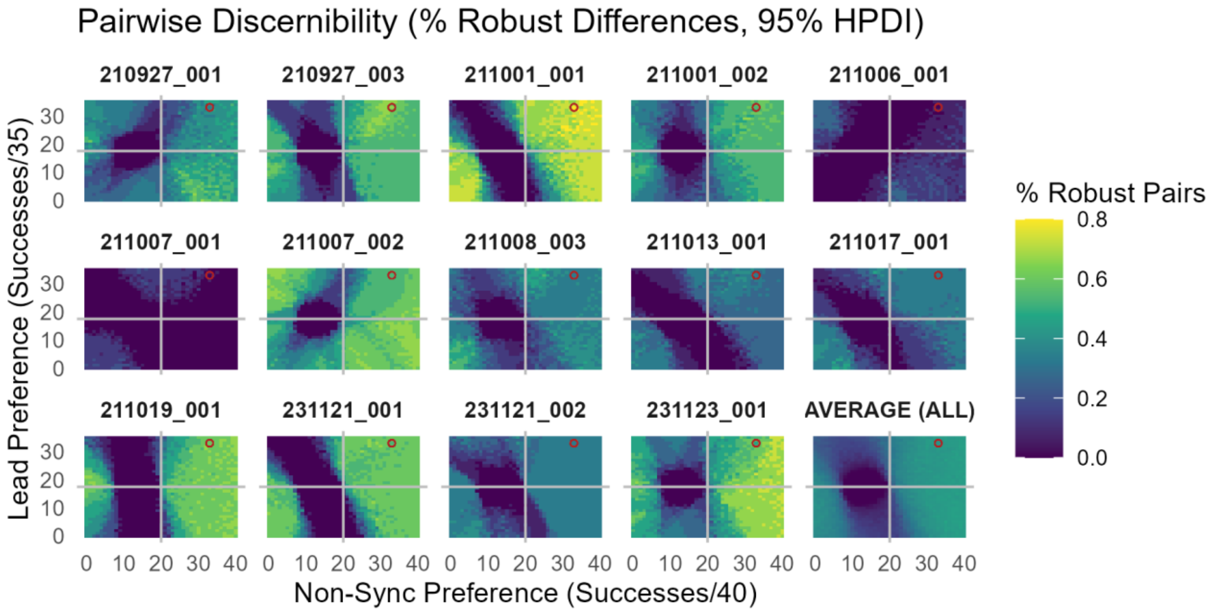


**FIG. S7**. the percentage of pairs within each chorus that differed robustly in $AC_{Synchrony}$ score. Red circle shows the true results of the previous female preference experiments used to calculate in-text results.

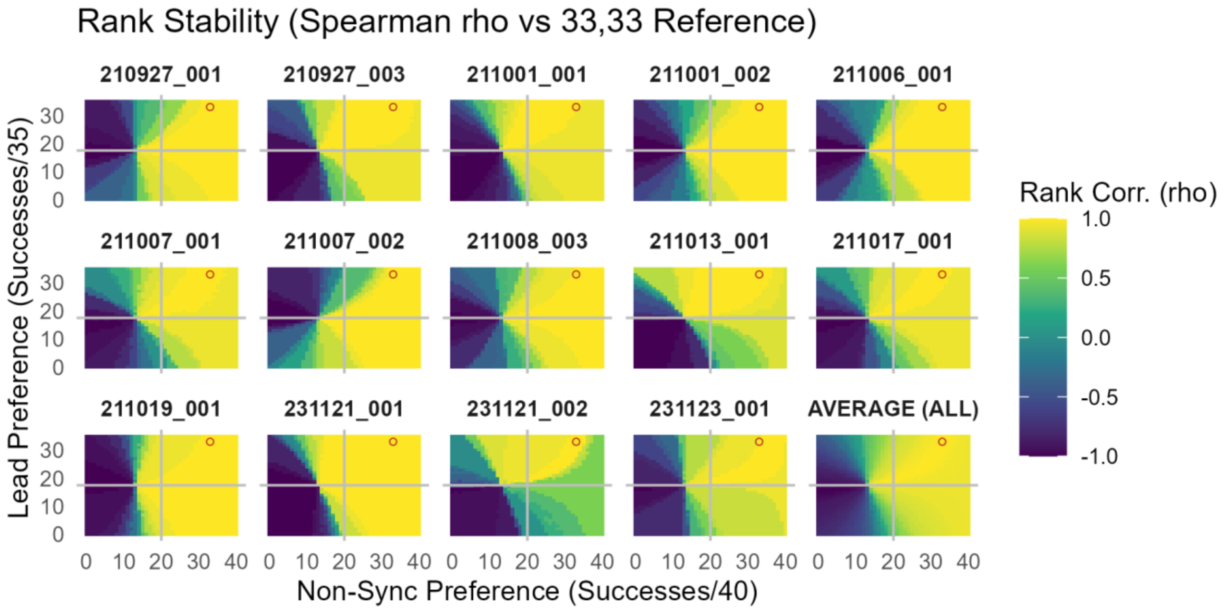


**FIG. S8**. the Spearman's rank correlation between males' mean $AC_{Synchrony}$ scores generated with true and alternate female preference values. Red circle shows the true results of the previous female preference experiments used to calculate in-text results.

**Supplementary Text S14. Selection Gradients Imposed on Sensorimotor Attributes by Synchrony Outcomes**

We wanted to know how the attractiveness consequences of engaging in synchrony (both synchrony rate and propensity to lead during synchrony) might be influencing selection on the sensorimotor attributes that drive these outcomes. Thus, we sought to reveal selection gradients imposed on intrinsic call period and leadership score (our proxy for effector delay duration) due to male synchrony engagement and the resulting attractiveness consequences. Our Bayesian bootstrapping analysis gave us a distribution of $AC_{Synchrony}$ scores per male that accounted for uncertainty in observed male behavioral tendencies and female preference estimates from previous experiments. To summarize each male's most probable attractiveness score, and the uncertainty surrounding this, we calculated the mean and standard deviation of his score distribution.

We then built Gaussian regression models with males' mean $AC_{Synchrony}$ score as the outcome variable and with measurement error weighted by the standard deviation around these means. This allowed us to propagate the uncertainty from our simulations into our statistical models. All models included random intercepts and random slopes for all main effects for chorus ID, as the strength of selection by female preferences is likely contingent on the exact social environment in which a male finds himself. We predicted that both intrinsic call period and leadership score would influence male attractiveness, and that they would interact as the mitigating effects of leadership score would be more pronounced for males with lower intrinsic call periods that synchronize more. Thus, we included leadership score, intrinsic call period and their interaction as fixed effects (syntax of this Full_Interaction_Model in Protocol 10 below). We also tested all sub-models.

We initially fit all models with 'sigma = TRUE' to model a global residual noise term on top of the individual uncertainty in male scores propagated through from our Bayesian bootstrapping simulations. However, this resulted in significant residual underdispersion across our models ($p = 0.024$) which manifested itself visually as severe vertical bunching in our residuals vs. fitted plots. This is likely because our Bayesian bootstrap pipeline (100,000 iterations) had already appropriately captured the uncertainty for individual male scores. Thus, adding this global residual variance parameter created redundant noise and artificially inflated the model's error bounds, rendering the true regression scatter underdispersed by comparison. Therefore, we opted to set 'sigma = FALSE'. This deactivated the global residual scale and conditioned the model's error structure entirely on the known, empirically derived bootstrap measurement errors (sd_attr). This completely resolved the underdispersion issues and resulted in well-fitting models.

- *Code for these models, model checks, model results and all visualizations here in RMarkdown file 'INDIVIDUAL_SYNCH_OUTCOMES.rmd'.*

**FULL_INTERACTION_MODEL**

```
Full_Interaction_Model <- brm(
  formula = mean_AC | se(sd_AC, sigma = FALSE) ~ Leadership_Score_std * Intrinsic_Call_Period_std +
                                                  (1 + Leadership_Score_std + Intrinsic_Call_Period_std |
Chorus_Name),
  data = data,
  family = gaussian(),
  prior = c(
    prior(normal(0, 1), class = "b"),
    prior(exponential(1), class = "sd")
  ),
  chains = 4, iter = 2000, cores = 4,
  control = list(
      adapt_delta = 0.999,   # For divergences
      max_treedepth = 20    # For treedepth warnings
    )
)
```

**Protocol 10**. Brms syntax for the full interaction model.

***Model Comparison Results***

| Predictor Model Structure | Δ elpd[1] | se (diff) | elpd_loo | se (elpd) | LOOIC |
|---|---|---|---|---|---|
| Additive_Model | 0.00 | 0.00 | 66.76 | 14.62 | -133.52 |
| Full_Interaction_Model | -1.23 | 0.90 | 65.53 | 14.82 | -131.06 |
| ICP_Only | -29.20 | 17.62 | 37.56 | 22.22 | -75.12 |
| Leadership_Rank_Only | -40.58 | 24.87 | 26.18 | 26.60 | -52.37 |
| Intercept_Only | -74.20 | 30.23 | -7.44 | 31.42 | 14.87 |

[1]The top-performing model is automatically anchored at Δ elpd = 0.00. While absolute intrinsic call period is the top model, note that deficits where Δ elpd / se < 2.00 represent highly competitive alternative structures.

**Table S18**. LOO-IC model comparison table for comparing the predictive power of the full interaction model and all sub-models.

*The Additive_Model* (just the main effects with no interaction term) was the top performing model overall, though it did not differ substantially from the *Full_Interaction_Model*. Both the *Additive_Model* and the *Full_Interaction_Model* were substantially superior to the intercept-only model in predictive performance ($\Delta elpd/se_{diff} > 2$).

***Results of Full_Interaction_Model:***

| Parameter | Median | 95% HDI Lower | 95% HDI Upper | pd | R-hat | ESS |
|---|---|---|---|---|---|---|
| ***Population-Level (Fixed) Effects*** | | | | | | |
| b_Intercept | 0.455 | 0.435 | 0.474 | 1.000 | 1.002 | 2,721 |
| b_Leadership_Score_std | 0.029 | 0.002 | 0.060 | 0.982 | 1.002 | 1,968 |
| b_Intrinsic_Call_Period_std | -0.026 | -0.051 | -0.001 | 0.981 | 1.001 | 2,643 |
| b_Leadership_Score_std:Intrinsic_Call_Period_std | 0.006 | -0.015 | 0.024 | 0.719 | 1.000 | 3,728 |
| ***Group-Level (Random) Effects (SD)*** | | | | | | |
| sd_Chorus_Name__Intercept | 0.028 | 0.012 | 0.050 | 1.000 | 1.001 | 2,190 |
| sd_Chorus_Name__Leadership_Score_std | 0.047 | 0.024 | 0.077 | 1.000 | 1.002 | 1,828 |
| sd_Chorus_Name__Intrinsic_Call_Period_std | 0.036 | 0.018 | 0.060 | 1.000 | 1.001 | 2,144 |

**Table S19.** Medians and upper and lower 95% *HDI*s of posterior distributions of fixed effects, random effects, and smooth terms. Also shown are Probability of Direction (*pd*), R-hat, and effective sample size.

***Fixed Effects Visualizations for of Full_Interaction_Model:***

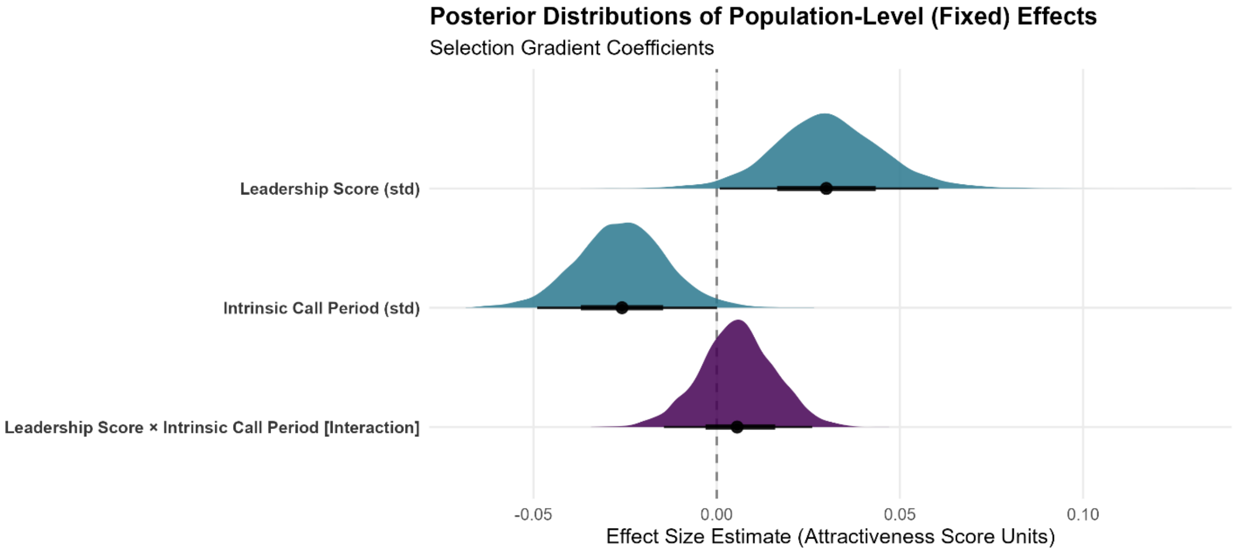


**FIG. S9**. Posterior distributions of population-level (fixed) effects on $AC_{Synchrony}$ scores (*Full_Interaction_Model*). Distributions are posterior parameter estimates derived from the Bayesian LMM. Density curves depict the full posterior probability density, points denote the posterior medians, and horizontal bars indicate the nested 66% and 95% Highest Density Intervals (*HDI*s).

***Random Effects Visualization for of Full_Interaction_Model:***

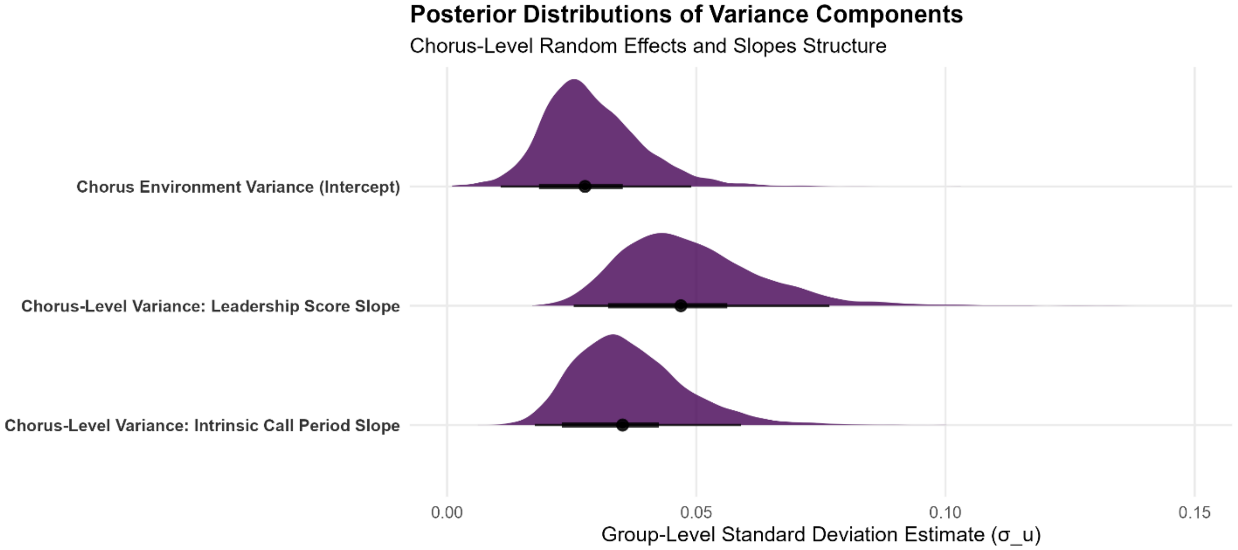


**FIG. S10**. Posterior distributions of group-level (random) effects and smooth term variances for $AC_{Synchrony}$ scores (*Full_Interaction_Model*). Distributions represent posterior parameter estimates (on the standard deviation scale) derived from the Bayesian LMM. Density curves depict the full posterior probability density, points denote the posterior medians, and horizontal bars indicate the nested 66% and 95% Highest Density Intervals (*HDI*s). These variance components quantify the structural variation in baseline probability and slopes across sampling units.

As expected, intrinsic call period had a robust negative effect on $AC_{Synchrony}$ scores (B-LMM: $\beta_{median}$ = -0.03, 95% *HDI* = [-0.05, -0.001], *pd* = **98%**), while leadership score had a robust positive effect (B-LMM: $\beta_{median}$ = 0.03, 95% *HDI* = [0.001, 0.06], *pd* = **98%**) (Fig. 3D, in-text; Supplementary Text S11). Their interaction was non-robust (B-LMM: $\beta_{median}$ = 0.01, 95% *HDI* = [-0.02, 0.03], *pd* = 72%). Interestingly, the effects of intrinsic call period and leadership score were almost perfectly symmetrical, despite three major underlying asymmetries in the system: **1)** their effects on their salient behavioral outcomes differed in magnitude (effect of intrinsic call period on synchrony rate: B-GLMM $\beta_{median}$ = 0.10, 95% *HDI* = [0.05, 0.15], *pd* = **100%**; effect of leadership score on follow rate: B-GLMM $\beta_{median}$ = -0.18, 95% *HDI* = [-0.22, -0.14], *pd* = **100%**), **2)** female preferences for their corresponding behavioral outcomes differed in magnitude (33/40 females choosing non-synchronous calls over synchronous calls (*4*); 33/35 females choosing leading calls over following calls during synchrony (*15*)), and **3)** both predictor variables were standardized separately relative to their own mean and standard deviation [(x-mean(x))/SD(x)].

Importantly, this symmetry is not a foregone conclusion or an artifact of our analytical pipeline. Changing female preference values during our simulation can yield non-symmetrical parameter estimate distributions that differ in both the magnitude of estimates and their robustness (Fig. S11). Thus, the fact that our experimentally derived female preferences interact with the observed variation in the corresponding male behavioral outcomes and their sensorimotor traits to yield selection of equivalent strength is simply a fascinating emergent property of this system.

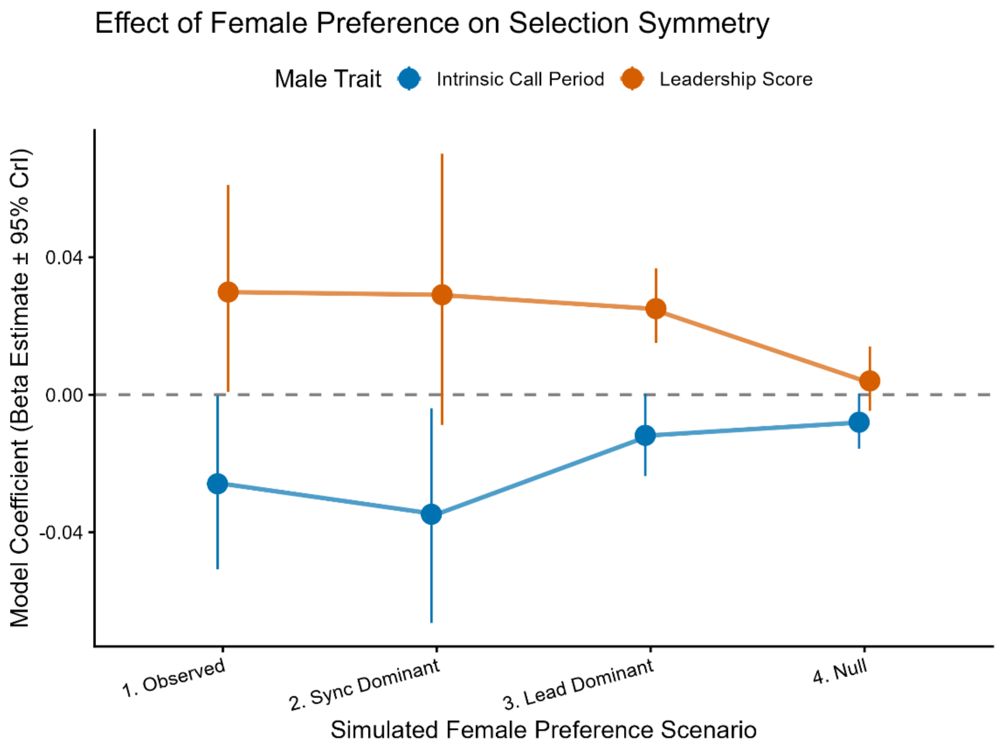


**FIG. S11**. Posterior median and 95% *HDI*s of parameter estimates from versions of the *Full_Interaction_Model,* when predicting male $AC_{Synchrony}$ scores simulated under different female preference strengths for non-synchronous calls and leading calls. The four simulation scenarios include: **Observed** (33/40 females chose non-synchronous; 33/35 females chose leading); **Sync Dominant** (38/40 non-synchronous; 18/35 leading); **Lead Dominant** (22/40 non-synchronous; 33/35 leading); and **Null** (20/40 non-synchronous; 17/35 leading). When female preferences are altered, the parameter estimates do not symmetrically track one another in magnitude, certainty, or robustness (whether 95% *HDI* includes 0). That they are not completely decoupled from one another either is likely because intrinsic call period and leadership score are somewhat associated (Supplementary Text S12), and therefore so are the behavioral outcomes that they lead to.
